# Terrestrial herbivory drives adaptive evolution in an aquatic community via indirect effects

**DOI:** 10.1101/2024.09.27.615417

**Authors:** Martin Schäfer, Antonino Malacrinò, Christoph Walcher, Piet Spaak, Marie Serwaty- Sárazová, Silvana Käser, Thea Bulas, Christine Dambone-Bösch, Eric Dexter, Jürgen Hottinger, Laura Böttner, Christoph Vorburger, Dieter Ebert, Shuqing Xu

## Abstract

Indirect ecological effects, which occur when the impact of one species on another is mediated by a third species or the shared environment, are ubiquitous in nature. Given the complexity of natural systems, indirect ecological effects were thought to be important in driving eco-evolutionary processes across community boundaries. However, we know remarkably little about such effects. Here we show that indirect effects of terrestrial insect (aphids) herbivory on macrophytes (duckweed) drives adaptive evolution of water fleas (*Daphnia*) in large outdoor aquatic mesocosms. Aphid herbivory reduced macrophyte growth and increased the abundance of phytoplankton, which in turn increased the abundance of *Daphnia*. Whole genome pool sequencing and phenotypic assays revealed an impact on the genetic compositions of the *Daphnia* populations and transplant experiments indicated that these evolutionary changes were adaptive. Furthermore, these changes in the aquatic community altered the interactions of the aphids and the macrophytes. These results demonstrate that indirect ecological effects can shape eco-evolutionary interactions between different communities.

## Introduction

Indirect ecological interactions are ubiquitous in natural ecosystems, where organisms live and evolve (*1*). Understanding whether and how indirect interactions shape evolutionary processes is a major challenge because the effects we observe when studying pairwise species interactions often cannot be extrapolated to more complex natural communities. Theoretical models suggest that indirect effects can drive coevolution in mutualistic networks (*2, 3*) and that cascading trophic interactions can influence species fitness across community boundaries (*4*). However, examples directly demonstrating that indirect interactions can shape evolutionary processes between natural communities are largely lacking. Furthermore, evolution can take place over ecological time scales of very few generations (*5, 6*), which can, in turn, affect ecological interactions. Given the complexity of natural ecosystems, it is important to extend our understanding of adaptive processes and eco-evolutionary dynamics across communities.

In natural communities, antagonistic interactions, such as plant-herbivore interactions and interspecific competition, are widespread and can have profound effects on the fitness of each species (*7–11*). In many shallow freshwater ecosystems, such as rivers, ditches, and ponds, macrophytes grow on the water surface and are attacked by both terrestrial and aquatic herbivores. In the water, macrophytes compete with phytoplankton for dissolved nutrients and light (*12*), and phytoplankton is a food resource of zooplankton. In such ecosystems, macrophytes can act as keystone species that mediate indirect interactions between terrestrial and aquatic communities (Fig. 1A). Here, we tested whether changes in the intensity of terrestrial herbivory on macrophytes can drive the evolution of species in an aquatic community via indirect effects, and whether these changes in the aquatic community can feedback on interactions between macrophytes and terrestrial insect herbivores. To this end, we established a replicated semi-natural aquatic community using outdoor experimental ponds (Fig. 1B and 1C). We grew the macrophyte *Spirodela polyrhiza* (giant duckweed) in the experimental ponds and manipulated the intensity of herbivory by waterlily aphids (*Rhopalosiphum nymphaeae*) on the duckweed. We then measured the ecosystem changes in the aquatic community and assessed the evolutionary responses of *Daphnia magna*, a key consumer of phytoplankton. By performing transplant experiments, we quantified the consequences of biotic and abiotic changes in the aquatic community on the fitness of aphids and duckweed, and assessed the eco-evolutionary dynamics mediated by indirect interactions.

**Fig. 1.**
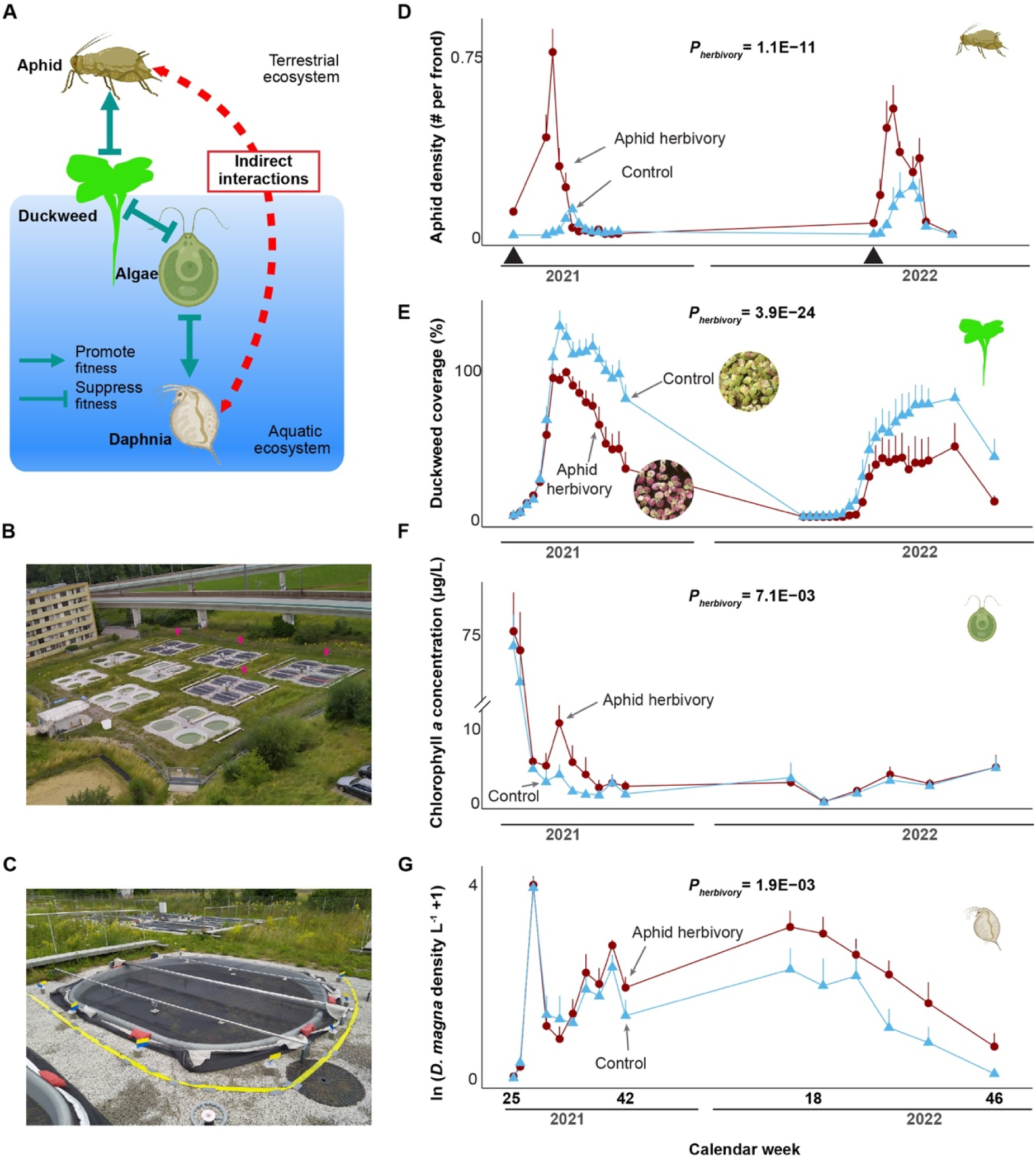
Aphid herbivory suppressed duckweed population growth and increased *Daphnia magna* abundance. A: a simplified species interaction network in the community. B: an overview of the outdoor experimental ponds. Arrows indicate the 4 x 4 ponds used for the experiments. C: a close view of an experimental pond, covered with a net to reduce aphid migration. D-G: the 2-year population dynamics of the aphids, duckweed, total phytoplankton (estimated by total chlorophyll a), and *D. magna*. The calendar week is shown on the x-axis. Aphid density (D), percentage of surface area coverage of the duckweed (E), chlorophyll-a concentration (F), and *D. magna* population density (in log scale) (G) are shown in the y-axis. The black arrows on the x-axis of panel D refer to the time when aphids were added to the herbivory ponds (experimental treatment). Dark red and light blue colors refer to herbivory and control treatment. Representative pictures of the duckweeds at calendar week 32 from the two treatments are shown in panel E, illustrating the marked color difference. Overview pictures of the ponds can be found in Figs. S4-S12. P-values were estimated using linear mixed-effects models with time and pond block as random factors. Panel A and species icons are created with BioRender.com.

## Results

### Aphid herbivory reduced duckweed growth and altered the aquatic community

To assess the effects of aphid herbivory on the aquatic community, we quantified the aphid population size, the duckweed growth, and the plankton communities over two consecutive years in the outdoor experimental ponds (2021 and 2022; Fig. S1). The aphid and duckweed populations grew exponentially in summer and declined to zero in winter (Fig. 1D, 1E, and Fig. S2A), when the duckweed formed resting stages (turions) that sank to the bottom of the ponds. We reintroduced aphids into the community in spring of the second year, 2022. In late summer (between calendar weeks 30 and 36), aphid herbivory reduced the duckweed population size by ∼30.7% and 50.1% in 2021 and 2022, respectively (Fig. 1E). In the first year, the herbivory-driven reduction of the plant population started shortly before the peak of the aphid population (calendar week 30). The reduced plant growth effect by aphid herbivory was still visible in spring 2022, before we reintroduced aphids into the ponds, suggesting that herbivory has a long-lasting influence on duckweed growth. In addition to reducing the growth rate and population size of the duckweed, aphid herbivory also increased the content of anthocyanins in the duckweed, such as cyanidin-3-O-glycoside and cyanidin-3-O-(6-O-malonyl-beta-glucoside), likely contributing to the color differences of the duckweed visible during late summer, with predominantly greenish leaves in the controls, and reddish leaves in the plants exposed to aphid herbivory (Fig. 1E small pictures, and Fig. S3).

Aphid herbivory on duckweed led to a trophic cascade effect in the aquatic community. In 2021, the reduced duckweed growth was shortly later followed by an increase in phytoplankton abundance (quantified using the chlorophyll-a concentration in the water as a proxy) and diversity between calendar weeks 30 and 35 (Fig. 1F and Fig. S2F). Shortly after that, we observed an increase in the population size of *D. magna*, the main consumer of phytoplankton. The increased abundance of *D. magna* in the herbivory ponds is the likely cause for the subsequent decline of the phytoplankton in this treatment, which remained low until the end of the experiment, while the *Daphnia* populations remained high until the end of the experiment in 2022 (Fig. 1G). From week 42 of the first year until the end of the experiment (more than a year), the average *Daphnia* population size in the herbivory treatment was approximately doubled in comparison to the control treatment. In addition to changing in the phytoplankton community, aphid herbivory also altered the abiotic environment, such as increased total phosphorus concentration, total dissolved carbon, water temperature, and pH (Figs. S13 and S14). Together, these results show that aphid herbivory on duckweed altered both the biotic and abiotic environment of the aquatic community, which influenced the population size of *D. magna*.

### Aphid herbivory drove rapid adaptive evolution of *D. magna*

We then investigated whether the observed biotic and abiotic changes in the aquatic environment associated with aphid herbivory also drove the evolution of *D. magna*. To this end, we phenotypically typed genetic markers developed to test for the attachment (adhesion phenotype) of different isolates of the bacterial pathogen *Pasteuria ramosa*, a common and virulent parasite of *Daphnia* (*13–15*). This parasite was not present in our mesocosms, but the phenotypic markers nevertheless can be used to trace changes at polymorphic loci, as they are very fast to type, allowing big samples to be processed. As the attachment phenotype of *P. ramosa* to the cuticle of *D. magna* is genetically determined, changes in the frequencies of attachment phenotypes indicate evolutionary changes in the *D. magna* population. Attachment phenotypes were assessed with the attachment test (*13*) in the laboratory using *D. magna* taken from the initial population and from the evolved pond populations. We found that the attachment level to three of the five tested parasite isolates differed between *D. magna* populations that evolved in control and in herbivory ponds (Fig. 2). *Daphnia magna* populations evolved lower attachment probabilities to isolates P15 and P21 but higher attachment probability to the isolate P20 in the herbivory ponds compared to populations evolved in the control ponds (Fig. 2). The effects were consistent in 2021 and 2022. To verify the absence of *P. ramosa* in the communities at the end of the experiment, we searched the pool-seq data collected for each *D. magna* population for DNA traces of *P. ramosa*, and no fragments of the parasite DNA were discovered in any population, suggesting the changes in parasite resistances are likely due to genetic linkage, pleiotropy or a cost of resistance. Genetic linkage, i.e. the genes under selection being physically linked to the attachment genes, is of particular importance here, as *D. magna* reproduces mostly asexually and the attachment genes typed with the attachment test are known to be located in large polymorphic haplotypes (*14, 15*).

**Fig. 2.**
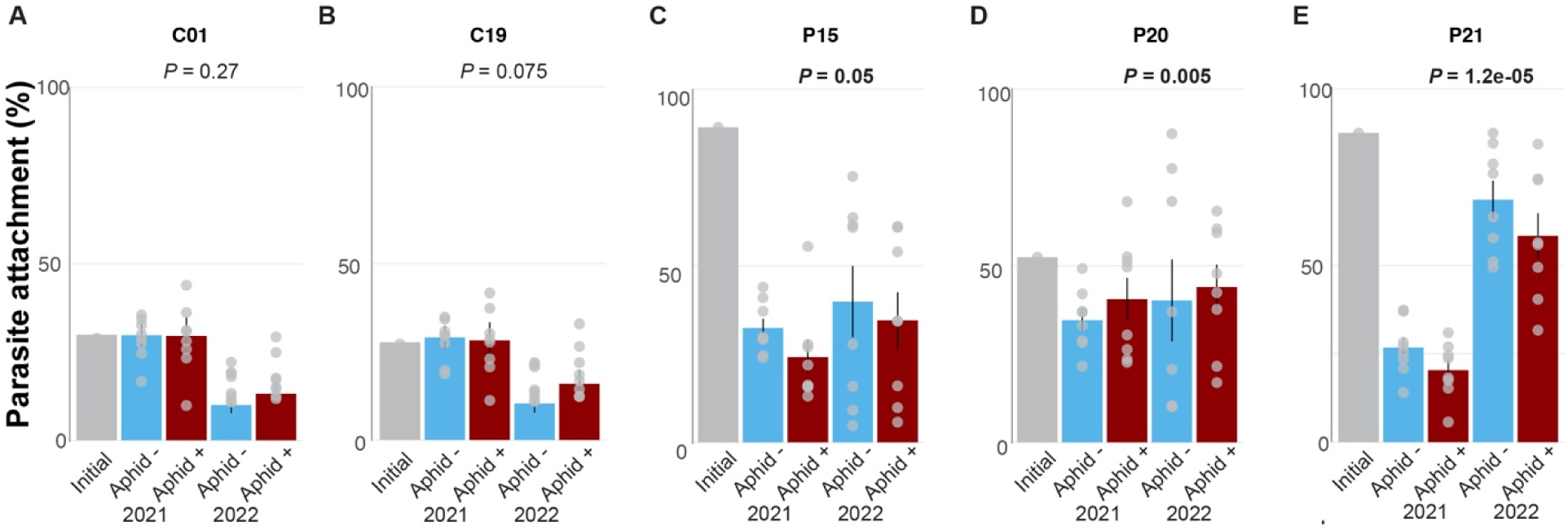
Aphid-herbivory changed *Daphnia magna* genetic composition. A-E refer to the changes of parasite attachment in the *D. magna* populations. Parasite attachment (quantified as the ability of the parasite *P. ramosa* to attach to the host cuticle) was measured using five isolates of *P. ramosa* (*15*): C1, C19, P15, P20 and P21. *P*-values indicate whether aphid herbivory affected attachment to *D. magna* populations to each parasite strain. *P*-values were determined using linear mixed-effects models with time and pond block as random factors. Grey bars refer to the frequencies of attachment of the parasite strains in the initial population. Blue and dark red colors refer to control and aphid-herbivory treatments, respectively. Error bars depict one standard error. In each panel, the individual resistance level from each population is shown as light grey dots.

To investigate whether the evolution of *D. magna* driven by aphid herbivory is reflected on the genomic level, we sequenced the genomes of the evolved *D. magna* populations at three different time points (beginning, middle, and end of the experiment) using a pool-seq approach.

To identify genomic signatures of selection in *D. magna* that were imposed by aphid herbivory on the duckweed, we then compared the allele frequencies between the *D. magna* populations evolved in the control and the herbivory ponds. At the genome-wide level, the *F_ST_* between control and aphid herbivory populations increased from 0.039 (± 0.022) in 2021 to 0.097 (± 0.06) in 2022 (Figs. S15 and S16), indicating that aphid-herbivore on the duckweed imposes divergent selection continuously driving the differentiation between the *D. magna* populations from control and aphid herbivory ponds. Consistently, using the Cochran Mantel Haenszel test (*16, 17*), we found that the proportion of SNPs showing significant allele frequency differences between control and aphid herbivory ponds increased from 8.1 % in 2021 to 28.6 % in 2022 (Fig. S17). To identify the regions that were under divergent selection imposed by aphid herbivory during the two years, we identified loci that showed different allele frequencies between control and aphid herbivory ponds using the beta-binomial mixed-effects model (with year and pond block as random factors). In total, we found that the allele frequencies of 141 SNPs were significantly different (*P* < 0.05 after Bonferroni correction) between the two treatments (Fig. 3). Several of these significant SNPs are located at the beginning of chromosome 10, consistent with the selection analysis (Fig. S18). Among these 141 SNPs, 105 were in the genic region of 57 genes (Table S1).

**Fig. 3.**
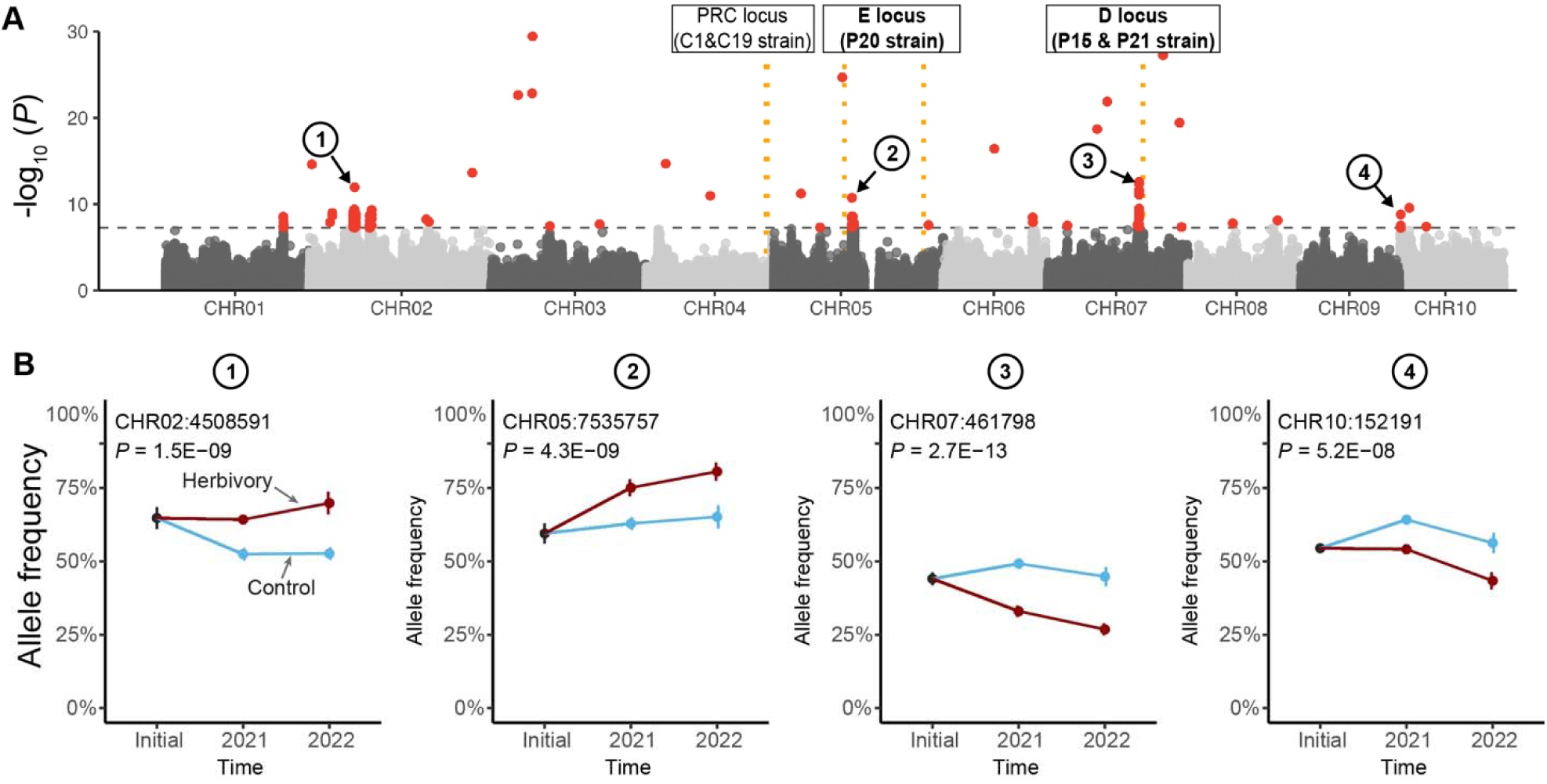
Aphid herbivory imposed genomic selection in *Daphnia magna*. A: Manhattan plot showing divergent selection imposed by aphid herbivory. Y-axis shows the –log_10_ P value of each SNP based on the beta-binomial mixed-effects model (with time and pond block as random factors). The dashed line refers to Bonferroni corrected *P* < 0.05 cutoff. Loci affecting the attachment of *P. ramosa* strains are shown in black boxes. PRC locus affects strains C1 and C19, the E locus affects strain P20, and the D locus affects strains P15 and P21. The *P. ramosa* strains that showed significant attachment phenotypes between control aphid herbivory ponds are highlighted in bold. B: plots showing allele frequency changes of four selected SNPs that are labeled in A. Blue and dark red colors refer to control and aphid-herbivory treatments, respectively. Error bar refers to standard error.

Several of the identified significant SNPs are located close to known loci affecting strain-specific parasite attachment and resistances (*15, 18*) (Fig. 3). Several significant SNPs are located within the large (nearly 5 MB) non-recombing haplotype that covers most of the right arm of Chromosome 5 and that determines the attachment of *P. ramosa* P20 strain to *D. magna*. Similarly, several significant SNPs are located on chromosome 7 near the D-locus that determines the attachment ability of *P. ramosa* strain P15 and P21 to *D. magna*. However, none of the significant SNPs were found near the *Pasteuria* resistance complex (PRC) locus on chromosome 4 (previously called ABCFG locus) that affects the attachment of *P. ramosa* strains C1 and C19. Thus, the changes in allele frequency near the parasite-attachment locus are highly consistent with the phenotypic changes observed in the populations (Fig. 2).

We further determined whether the observed evolutionary changes in *D. magna* are adaptive by performing transplant experiments in the second year of the experiment (2022) (Fig. S19). To this end, we collected *D. magna* individuals from each pond and placed them into both control and herbivory ponds, respectively, using PVC columns containing various cut-outs covered by mesh to allow for an exchange of water and phytoplankton (Fig. S19). We performed the same experiment once in spring (calendar week 23 to 25) and once in summer (calendar week 28 to 30). Both the season and pond community significantly affected the growth rate of *D. magna*. In calendar week 25, when *D. magna* grew rapidly in most of the ponds, the *D. magna* populations originating from aphid herbivory ponds exhibited significantly higher fitness when they were placed inside the aphid herbivory ponds compared to control ponds (*P* = 0.004, *F*-test, Fig. 4). The *D. magna* evolved in the control ponds had similar fitness in both the herbivore and control ponds (*P* = 0.73, *F*-test, Fig. 4). In calendar week 30, *D. magna* population size did reproduce anymore in most of the ponds and thus their growth rates were similar between the treatments. Together, these data suggest that the evolution of *D. magna* populations in the aphid herbivory ponds reflects an adaptive response to this environment.

**Fig. 4.**
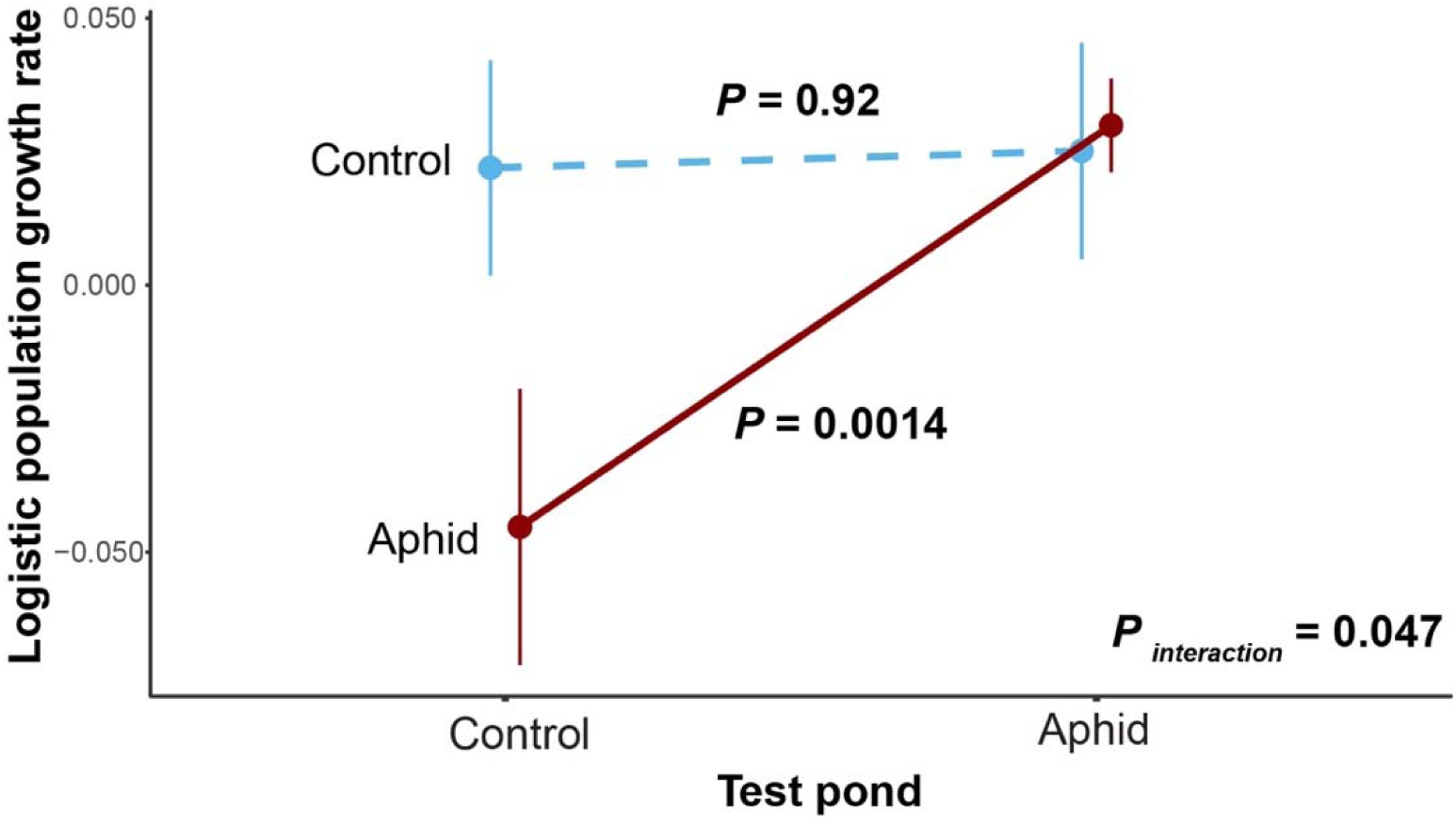
Evolution of *D. magna* is likely adaptive. The x-axis shows the community environment in which the transplant experiment is conducted. The y-axis refers to the log-transformed growth rate of *D. magna*. Light blue and dark red color indicate the source community environment, i.e. the community in which the *Daphnia* evolved. *P*-values refer to the effects of the testing community. *P_interaction_* refers to the interaction effects between testing community and evolution. Mean and standard error are shown.

### Changes in the aquatic community altered the aphid-duckweed interaction

The changes mediated by aphid herbivory in the aquatic community, in turn, might also alter the interactions between the aphids and the duckweed and thus lead to eco-evolutionary feedback. To test this hypothesis, we quantified the growth rate and aphid resistance of four duckweed genotypes by growing aphids and duckweed inside the evolved aquatic environments (Fig. S20). We performed the experiments twice in 2022, once in late spring (from calendar week 20 to 25) and once in summer (from calendar week 25 to 30). While the four duckweed genotypes had different growth rates (*P* < 0.001, *F*-test, Fig. 5), the changes in the aquatic communities caused by aphid-herbivory increased the duckweed growth rate in the absence of aphids (*P* = 0.036, *F*-test, Fig. 5). The magnitude of the community effects was however only 11% of the direct suppression effects from the aphid herbivory on the growth rate (e.g., at calendar week 30). The interaction effect between the duckweed genotype and community was, however, non-significant (*P* = 0.24), likely due to the small sample size. It is worth noticing that the genotype (SP102) that we used to establish the initial population appears to show the highest growth rate increase in the aphid-herbivory environment. This indicates that the environmental feedback on the duckweed might be genotype-specific, which in turn might shape the evolutionary processes of the duckweed-aphid interactions. Among the four duckweed genotypes, aphid growth rates differed (*P* < 0.001, *F*-test), suggesting that host plants varied in their defense or nutritional value. Interestingly, aphids grew faster in the aphid-herbivory ponds than in the control ponds (*P* = 0.003, *F*-test), regardless of the tested plant genotype, suggesting that aphid-herbivory induced community changes had positive feedback effects on aphids.

**Fig. 5.**
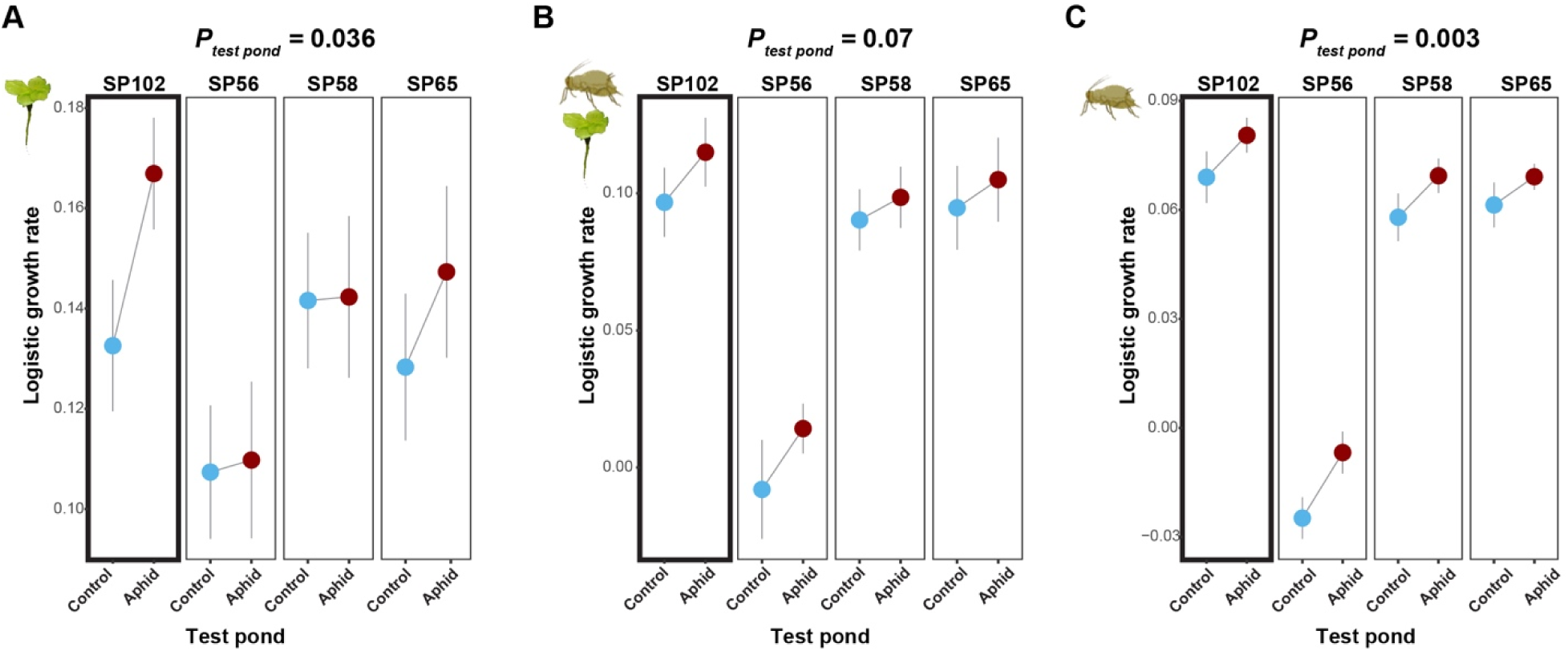
Evolution of the aquatic community altered the aphid-duckweed interactions. The x-axis shows the community environment in which the transplant experiment was conducted. The y-axis refers to the logistic growth rate of the duckweed without (A) or with aphid herbivory (B). In panel C, the y-axis refers to the logistic growth rate of the aphid. The average values from the two repeated experiments are shown. Light blue and dark red colors indicate the community environment where the plant and aphid fitness were quantified. *P*-values refer to the effects of the testing environment with experiment time as a random factor. All three parameters were measured in four duckweed genotypes that were collected in Europe (SP102, SP56, SP58, and SP65). Duckweed genotype SP102 was used in the mesocosm experiment (marked with a thick line box). Mean and standard error are shown.

## Discussion

In natural communities, indirect ecological effects can play an important role in shaping community structure and affecting organisms’ fitness (*2–4*). Here, by experimentally manipulating herbivory by a terrestrial insect on aquatic plants and measuring evolutionary responses in the planktonic crustacean *D. magna*, we demonstrated that indirect ecological interactions can drive adaptive evolution in a natural multispecies community.

The observed evolutionary changes in the large and genetically diverse *D. magna* populations are likely due to a combination of changes in the quality and quantity of their planktonic food, as well as abiotic changes in the environment caused by aphid herbivory on duckweed, such as different light and temperature conditions caused by the difference in the amount of shading by the duckweed. The phytoplankton abundance and diversity reacted strongly to the herbivory treatment in the first year (Fig. S2), likely due to the increased light and nutrient availability, and higher temperature in aquatic environments. Such changes can affect the growth of *D. magna* in a genotype-dependent manner (*19, 20*), driving divergent evolution of the *D. magna* populations in the two treatments. In addition to changes in the phytoplankton community, the herbivory treatment also altered the abiotic environment, such as temperature and pH (Fig. S14), two factors that are known to affect *D. magna* growth and reproduction (*21, 22*). Accordingly, transplant experiments suggested that *D. magna* evolved in the aphid herbivory ponds had lower fitness when they were placed in the control ponds. However, *D. magna* that evolved in the control community showed similar fitness in both communities, indicating an asymmetry in the adaptation to the herbivory-induced community changes.

Our genomic analysis revealed that the aphid herbivory-imposed selection shaped the genomic changes in the *D. magna* populations (Figure 3). The genome-wide population differences between *D. magna* populations evolved to become stronger over time between control and herbivory ponds (2.4-fold higher in 2022 than in 2021), indicating the cumulative effect of indirect selection imposed by the treatment. Because the levels of sexual reproduction and recombination in *D. magna* were limited during our experimental phase, we expected that a large proportion of the allele frequency changes in the populations would be affected by clonal selection, during which the entire genome is in genetic linkage. Consistently, the phenotype frequencies of the parasite attachment genes, which we used as convenient genetic markers, diverged among the treatments. The relevant parasite was not present in our mesocosms. All but one of these attachment genes are in large non-recombining regions in the *D. magna* genome (*14, 15*), suggesting that linked genes may have been affected by selection. These haplotype blocks are known to evolve very fast in *D. magna* populations (*23*). Thus, the change in parasite attachment phenotype frequency might have been a side-effect of selection on nearby genes. This would indicate that if parasites had been present, their interaction with the host would have been impacted by the herbivory treatment as well. Alternatively, selection by the parasite on the host (*23*) might influence the response of *D. magna* to the herbivory treatment, potentially influencing how other components of the system respond. The evolutionary changes observed are consistent among replicates in the two treatments. This and the fact that the *Daphnia* population was at no moment in time smaller than 1,200 individuals, exclude genetic drift as an explanation for the observed changes.

In our experiment, we only introduced a single genotype of the aphid and the duckweed. Under natural conditions, populations of the aphids and the duckweed also harbor genetic variation and, therefore, can also evolve in response to changes in the community via direct and indirect interactions. This study, therefore, identifies a plausible path of how species living in terrestrial and aquatic communities can coevolve via eco-evolutionary feedback.

## Materials and Methods

### Establishing outdoor experimental ponds

In 2021, we established 16 outdoor experimental ponds at the Swiss Federal Institute of Aquatic Science and Technology (Eawag) in Dübendorf, Switzerland, using a similar setup as reported by Narwani et al. (*24*) (Figs. S1 and S21). Each pond has a size of approximately 4 m × 4 m, with a maximum depth of 1.5 m and a water volume of ∼15 m^3^. At the beginning of the experiment, each pond was cleaned with a high-pressure cleaner (Fig. S21D). Then, we added 220 L of leaf litter collected from a forest nearby (next to Wangnerwald, 47 24’33.5” N, 8 40’05.5” E) to each pond to provide nutrients and shelters for different aquatic organisms (Fig. S21E). After filling with tap water (calendar week 22), we inoculated the ponds with an initial plankton and microbe community collected from Lake Greifensee (47 20’59.592” N, 8 40’40.736” E), located near the experimental site (calendar week 24). In each pond, we added 27 L of lake water that was collected from 5-6 m depth after removing large organisms using a net with a pore size of 95µm. Additionally, we added 2.8 L concentrated plankton community (30-95 µm fraction), each collected from filtering approximately 1,750 L of lake water. The 30-95 µm plankton fraction was collected by sampling water columns from 0-10 m depth with a 30-µm mesh (Fig. S21F) and pouring the concentrate subsequently through a 95-µm mesh to exclude larger organisms. Since our focal plant species, *Spirodela polyrhiza*, is known to produce turions (resting stages) in low phosphate environments (*25*), we additionally added 40.8 g of KH_2_PO_4_ per pond (ca. 20 µM; calendar week 24).

### Establishing the aphid-duckweed-daphnia populations in the ponds

In calendar week 25 of 2021, we added into each pond approximately 1,300 *Daphnia magna* individuals from a mix of 122 genotypes (=clones)( Fig. S21J), additional algae (*Tetradesmus* spp. and *Nanochlorpsis* spp.) that were used to feed *D. magna* in the lab, approximately 5,000 *Spirodela polyrhiza* fronds from a single genotype (Figs. S21G and S22; genotype SP102, originally collected in Switzerland and pre-cultivated in the lab before the field experiment) covering approximately 1% of the water surface and 18 great pond snails (*Lymnaea stagnalis*, Fig. S21H). We introduced the snails in the ponds because our previous experiments suggested that the snails are important to stabilize the fresh water community by controlling the growth of macro-algae (*26*). All *D. magna* genotypes had originally been collected in Lake Aegelsee near Frauenfeld (47°33.48’N, 8°51.66’ E), Switzerland (*27*). These clones were produced from wild caught females in the years 2014 to 2019 and kept in the laboratory by clonal propagation.

All *D. magna* clones were propagated in 400-mL jars (6 jars per clone) with abundant food (green algae *Tetradesmus* sp.), at a 16:8 hours light:dark cycle and a temperature of 20 °C. When all clones were at their approximate carrying capacity, we mixed all clones into four 60-L tanks. The tanks were topped up to 45 L volume. This procedure was replicated 4 times. Subsamples were counted, resulting in an estimated population size of 7,300 *Daphnia* per 60-L tank. The following day, the four tanks were transported to the field site. The populations in the tanks were well mixed, and 20 subsamples of equal size (2 L volume) were collected from each of the four tanks, resulting in 20 samples of 8 L volume with an estimated 1,300 *D. magna* individuals. Sixteen sub-samples were used to inoculate the 16 experimental ponds. The initial inoculum for each mesocosm was large enough to minimize possible stochastic effects, influencing SNP and resistance phenotype frequencies. The remaining four samples were frozen in liquid nitrogen for later genomics analyses (see below), which were considered as samples from timepoint-0. From the timepoint-0 samples, we also collected 150 individuals of *D. magna* to estimate parasite attachment phenotype (see below).

In each pond, we introduced 200 waterlily aphids (Fig. S21I, *Rhopalosiphum nymphaeae*) from a single genotype into eight ponds while we kept the other eight ponds free from aphids as control. The aphid populations were developed from a single individual that was collected at the University of Münster campus (51°57’40.7” N, 7°36’55.8” E) in September 2020. The ponds were arranged in four blocks of four ponds, each containing two ponds with aphids and two control ponds. The populations of aphid, duckweed, and plankton communities were randomized.

To minimize aphid migration between ponds, we installed pond covers made from mosquito nets, and sticky flags and sticky tape were placed on the ground between ponds (Fig. S21C). Control ponds were regularly monitored for aphids. Aphids that accidentally invaded control ponds were manually removed as long as feasible.

The experiment was run for two years, i.e. two summers and the winter in between. The pond covers were removed over the winter season to avoid damage due to snowfall and reinstalled in March 2022 (calendar week 11). We reintroduced the aphids in calendar week 25 in 2022 to the same ponds that already received aphids the year before since the aphids do not overwinter within the ponds.

Due to an unexpected extinction (for unknown reasons) of pond snails in pond 6A in 2021, we removed this replicate from all analyses. We also excluded the 2022 data from ponds 1D and 3D from the aphid herbivory treatment because the aphid populations didn’t successfully establish in these two ponds in 2022.

### Monitoring population dynamics in the experimental ponds

Duckweed growth was assessed based on an estimation of the surface coverage of the ponds with duckweed. Values above 100% represent multi-layered growth (via close inspections). Additionally, at each timepoint overview pictures of the ponds were taken with a camera (GoPro Hero 7 and Hero 9) that was attached to a long stick. Aphid abundance was calculated based on close-up pictures that were taken with a mobile phone from 8 random positions of each pond (Fig. S21C). Within a subsection of four pictures per pond, all aphids and fronds were counted to calculate the number of aphids per frond.

We monitored the population dynamics of plankton organisms using a previously established protocol with minor modifications (*24*) (Fig. S23). Briefly, we sampled three vertical profiles of the water column using a Leibold-sampler, which consists of a 180 cm long PVC tube (5 cm diameter) that can be closed via a wire with a plug at the bottom (Figs. S21D and S21E). From each pond, we collected 10 L of water per sampling. From this, we took a 1 L subsample for nutrient analyses (Fig. S21F) and 1 L for chlorophyll-a and phytoplankton analyses (Fig. S21G). During this subsampling, zooplankton was prevented from leaving the beaker with a 150 µm mesh. The remaining water (with zooplankton corresponding to 10 L pond water) was filtered through a 150 µm mesh to collect the retained zooplankton (Fig. S21H).

We estimated the total phytoplankton abundance in the pond by measuring chlorophyll-*a* content in the water. To this end, each sample was filtered using a 25 mm GF/F filter on a 50 mL syringe. Samples were filtered until a noticeable backpressure occurred or the filter turned green. The volume of filtered water was noted for later normalization. Afterwards, excess water was removed from the filters by pushing another syringe filled with air through the filter. Filters were then removed from the support and shortly dried on a paper towel before they were placed in aluminium-covered tubes on ice and subsequently stored at –20°C until further processing. We extracted chlorophyll-a from the filters with 5 mL 90% EtOH after vortexing, sonication and overnight incubation at 4°C in the dark. Subsequently, the extracts were passed through 0.2 μm cellulose acetate syringe filters before they were analyzed on a Hitachi 2000UV/VIS Spectrophotometer LLG-uniSPEC4 based on the absorption at 750 nm and 665 nm and an external chlorophyll-a standard. The chlorophyll-a content was calculated using the following formula:

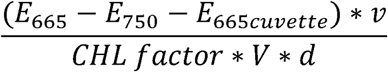

where *E_665_*refers to extinction at 665 nm, *E_750_* refers to extinction at 750 nm, *E_665cuvette_* refers to extinction at 665 nm of a cuvette filed with 90% EtOH, *v* refers to the volume of 90% EtOH, *CHL* factor refers to a correction factor (0.000082), *V* refers to the filtrated volume and *d* refers to the cuvette length (5cm).

To determine the taxonomic composition of phytoplankton, the water samples were preserved by adding each 5 mL of Lugol’s solution into 50 mL of pond water. Samples were stored at 4 °C until further processing. For analysis, subsamples were placed in a Utermöhl counting chamber for 24 hours. Chlorophyll-*a* concentrations were used as a guide for high densities samples, of which 3 mL were sedimented. Afterward, the tube was removed, a cover slide was placed on top of the counting chamber, and the sedimented plankton was analyzed under a Zeiss Axiovert 135 inverted microscope (*28*). To this end, phytoplankton were identified (mostly to genus level) and counted in 40 randomly chosen fields of view (out of 1,686) with 320X magnification and 40 fields of view (out of 6,900) with 640X magnification. If one taxon was highly abundant (>100 cells in 10 fields of view), we counted this taxon in only 10 fields of view.

The zooplankton collected from the 10 L of pond water via a 150-μm mesh was eluted from the mesh with 100 % EtOH each into a 50-mL Kautex bottle. Samples were stored at room temperature until further analyses. Samples were analyzed under a Leica M205 C stereomicroscope at 10-50X magnification, and counts were converted to individuals per litre of pond water. Adult Cladocera were identified at the species level, juvenile Cladocera and Copepods at the genus level, and other zooplankton at higher taxonomic levels.

### Monitoring nutrients and abiotic environment in the experimental ponds

Orthophosphate-phosphorus (PO_4_ ^3-^-P) was measured photometrically using an Agilent Cary 60 spectrophotometer after reaction to phosphorus molybdenum blue complex. Total phosphorous (P) was measured after decomposition by autoclaving at 121 °C with subsequent determination of PO_4_ ^3-^-P as above. Ammonium-nitrogen (NH ^+^-N) was measured photometrically using an Agilent Cary 60 spectrophotometer after reaction to a blue complex by Berthelot-reaction. Total organic carbon (TOC) was measured on a Shimadzu TOC-L CSH after catalytic combustion at 720°C with subsequent measurement of CO_2_ via an infrared detector.

The pH and oxygen concentrations of the pond water were regularly monitored using handheld sondes (Multi 3630 IDS with an FDO 925 oxygen sensor and a SenTix 940-3 pH sensor, all from WTW) (Fig. S23B). The dipping probes were placed 75-80 cm below the water surface. For the continuous analysis of the water temperature and light penetration into the pond water we deployed a logger in the middle of each pond 30-35 cm below the water surface (Fig. S23A). Six ponds (3 per treatment) were provided with a temperature logger (HOBO pendant temperature 64K, UA-001-64), and 10 ponds (5 per treatment) were provided with a temperature logger with a light sensor (HOBO Pendant Temperature+Light 64K, UA-002-64). The data were recorded every 15 minutes. The temperature was expressed as weekly average temperature. Light data (in LUX) were summed up for each day, and weekly averages of the daily sum of light were calculated.

### Assessing parasite attachment and genomic changes in the *D. magna* populations

At defined time points during the experiment, samples were collected to assess parasite attachment and genomic changes using a pool-seq approach. Samples at timepoint 0 were collected from the original inoculum, as described above. *Daphnia* population samples were collected from the experimental ponds. For this, we used handheld nets with a mesh width of about 0.2 mm. Care was taken to ensure that every depth and every area of the ponds were swept with the net. The samples were brought to a laboratory and were inspected using stereo microscopes. We sorted the samples to include only *D. magna*. From each sample, about 250 adult *D. magna* were collected and frozen with little water in liquid nitrogen. Sampling dates for the pool-seq samples were June 22nd, 2021 (timepoint 0, calendar week 25), September 22nd 2022 (calendar week 38), and July 28th, 2022 (calendar week 30). Samples for the assessment of parasite attachment phenotypes were transported in 2-L bottles to the University of Basel laboratory, where 100 animals from each pond were placed individually in a 100-mL jar. Animals were tested for parasite resistance as described below. Sampling dates roughly coincided with the sampling dates for the pool-seq samples but were more spread out, because the workload associated with testing 16 times 100 animals was too high to be performed in a short time interval. Therefore, samples were taken across a time period of about 4 weeks (from September 22nd to October 16th, 2021 and from July 3rd to July 28th, 2022). Within each sampling campaign, samples were randomized (treatment and blocks were exactly balanced), and the assessment of attachment phenotypes was blinded with regard to the treatment a sample came from.

For each clone we assessed the attachment of five isolates of the common bacterial pathogen, *Pasteuria ramosa*, using the attachment assay (*29*). In short, fluorescently labeled parasite endospores were exposed to *D. magna* individuals. After 20 to 60 minutes, animals were assayed with a fluorescence microscope searching for endospores attached to the foregut or the hindgut. The ability of the parasite to attach to the gut epithelium is a prerequisite for infection and correlates very strongly with successful infection (*29*). The loci determining attachment are mendelian, but interact epistatically with each other (*14, 15, 23*). The diversity of our panel of *D. magna* clones was enriched for rare attachment phenotypes.

For the pool-seq samples, we extracted DNA from pooled individuals from each pond using the CTAB method. The total DNA was sequenced with 150 bp paired-end reads at ∼480X coverage for each sample using an Illumina NovaSeq 6000 instrument by Novogene in Beijing, China. Raw data was quality-checked and trimmed using TrimGalore v0.6.1 (*30*), and reads were mapped towards the *D. magna* reference genome (*31*) using BWA (*32*) and SAMtools (*33*).

Signatures of selection in *D. magna* populations from both herbivory and control ponds were detected using CLEAR based on time-series data (*34*). We then used SAMtools (*33*), PoPoolation2 (*35*) and poolfstat (*36*) to estimate genome-wide *F_ST_* between and within treatments and sample time points, with a minimum read coverage of 20 and a maximum coverage of 400 as the cutoff. The principle component analysis was performed using the R-package “pcadapt” (*37*). The heatmap showing the similarity among each pool was created using a pairwise *F_ST_* matrix estimated by poolfstat. To identify allele frequency changes for each SNP within each time point, we used Cochran Mantel Haenszel (CMH) test (*38, 39*). We used Bonferroni’s corrected *P*-value of 0.05 as the cutoff to identify significantly differentiated genomic regions. To identify genomic regions that were under selection during the two years, we used the beta-binomial mixed-effects model (from R-package glmmTMB v1.1.9 (*40*)) with sampling time and pond block as random factors. The genomic position of the resistance locus was identified using the sequence from Fredericksen et al. (*18*).

We used the pool-seq data to test for the presence of *P. ramosa* in our samples. Using SAMtools, we selected the sequences that were not mapped to the reference genome of *D. magna*. These data were then used as input for Kraken2 (*41*) to characterize the *Daphnia*-associated microbial community using a custom reference dataset that included the bacterial and fungal RefSeqs from NCBI and, additionally, the reference genome of *P. ramosa* (*42*). This analysis did not identify any sequence of the parasite in any of our samples.

At the genome-wide level, the populations in the ponds differentiated from the initial population over time (as measured by *F_ST_*) (Figs. S15 and S16). We then estimated genome-wide selection based on the time-series allele frequency changes. We found strong signatures of selection on chromosome 4 in both control and aphid herbivory populations (Fig. S18). The region on chromosome 4 overlapped with the known parasite-attachment genetic region (*18*), suggesting this region is under strong selection in the pond environment, a pattern that is consistent with the observed rapid changes in parasite attachment phenotypes in 2021 and 2022 in comparison to the initial population (Fig. 2). Furthermore, we found significant signatures of selection at the beginning of chromosome 10 only in the herbivory ponds (Fig S17), but not in control ponds, indicating divergent selection on the *D. magna* between herbivory and control ponds.

### Transplant experiments

To assess the effects of the aquatic community on the *D. magna* growth, we used a 140 cm long vertical PVC tube (5 cm diameter, Fig. S19). The tube was closed at the bottom and contained 4 equally distributed openings (12 × 6 cm) that were covered with a 150 µm mesh to allow for water and nutrient exchange. At the beginning of the transplant experiments, we added 50 similar-sized *D. magna* females (without eggs) originating from the ponds into each tube. The *D. magna* were sorted using a stereomicroscope. The *D. magna* were either kept in the same pond, moved to another pond from the same treatment or moved to another pond with different treatment. At the end of the experiment, the *D. magna* inside each tube were collected on a 1 mm mesh, from which they were kept in 100% EtOH. Samples were stored in a 50 mL Kautex bottle at room temperature until further analyses. We counted *D. magna* individuals using a Leica M205 C stereomicroscope. We performed the experiments twice. The first experiment lasted for 13 days (between calendar week 23 and 25), and the second experiment lasted for 12 days (between calendar week 28 and 30). The logistic growth rate was calculated as log(N_end_) – log(N_start_), where N_end_ and N_start_, refer to the number of *D. magna* at the end and beginning of experiments, respectively.

To assess the effects of the aquatic environment on the fitness of the duckweed and aphid, we performed growth assays using the floating boxes (Fig. S20), which allowed for the exchange of water and plankton while preventing the escape of the duckweed and aphids via mesh. The floating boxes were constructed using plastic boxes with lids (Eurobox, Auer Packaging, inside measures: w × d × h 17 cm × 12 cm × 11.5 cm). We cut holes in the bottom (15.3 cm × 10.3 cm) and lid (16.8 cm × 11.8 cm) and covered them with steel meshes (bottom: mesh size 0.63 mm, wire diameter 0.22 mm, material 1.4301 steel; lid: mesh size 0.14 mm, wire diameter 0.112 mm, material 1.4301 steel). Each box was fitted with a styrodur frame on the outside to make it float on the pond surface.

We performed the growth assays using four duckweed genotypes (SP102, SP56, SP58 and SP65) that were originally collected in different areas in Europe (*43*). At the beginning of the experiment, we used approximately 20 duckweed individuals (fronds) from one of those genotypes for each floating box. For each pond and each genotype, we used two boxes: in one, we added five aphids, while the other one was left without aphids (control). Approximately two weeks later, we collected both duckweeds and aphids and froze them in liquid nitrogen. The dry weight was measured after freeze-drying. To estimate the number of individuals, we collected and counted up to 100 duckweed individuals for each box and measured their dry weight. Then, dry weight per frond was calculated and used to estimate the total number of duckweed individuals for each box based on their total dry weight. To estimate the number of aphids in each box, we also determined the number of aphids per frond from those up to 100 fronds and multiplied it by the total number of fronds per box. The logistic growth rate was calculated as log(N_end_) – log(N_start_), where N_end_ and N_start_ refer to the number of duckweed or aphids at the end and beginning of experiments, respectively. The resistance of duckweed was estimated by calculating the differences in growth rate between control and aphid-treated boxes for each genotype inside each pond.

### Anthocyanin analysis in the duckweed populations

Samples from the duckweed populations in each pond were collected, cleaned from aphids and other large contaminants, washed with fresh tap water, dried using paper towels, and shock frozen in liquid nitrogen. Samples were kept until further processing at –20 °C. Anthocyanins were analyzed by LC-MS and HPLC-PDA, similar as described by Malacrinò et al. (*43*). In brief, approximately 10 mg of fine ground, freeze-dried plant tissue was extracted with acidified methanol. Fresh weight of was estimated based on the fresh weight to dry weight ratio of each sample. Cyanidin-3-O-glucoside was analyzed by LC-MS. Before analysis the sample extracts were diluted 1:100 in an aqueous mix of isotope-labeled amino acids (algal amino acid mixture-^13^C-^15^N; Sigma-Aldrich). Chromatographic separation was done on a Shimadzu Nexera X3 equipped with an Agilent 1290 infinity II inline filter (0.3 µm) and a ZORBAX RRHD Eclipse XDB-C18 column (3×50 mm, 1.8 µm; Agilent Technologies). Separation was done in gradient mode with 0.05 % formic acid (Fisher Chemical), 0.1% acetonitrile (Fisher Chemical) in water as Solvent A and methanol (Fisher Chemical) as Solvent B, with the column oven set to 42 °C and a flow rate of 500 µL/min using following solvent settings (Time [min]/B[%]): 0.0/2, 1.5/2, 3.5/100, 4.5/100, 5.0/2, 6.0/2. The LC-system was coupled to a Shimadzu LCMS-8060 mass spectrometer equipped with an ESI source, which was operated in positive ionization mode and the following settings: Nebulizing Gas Flow: 3 L/min; Heating Gas Flow: 10 L/min; Drying Gas Flow: 10 L/min; Interface Temperature: 300 °C; DL Temperature 250 °C; Heat Block Temperature: 400 °C; CID Gas: 270 kPa; Q1 Resolution: Unit; Q3 Resolution: Unit. Analysis was done in the multi-reaction-monitoring (MRM) modus using following precursor to product ion transitions: Cyanidin-3-O-glucoside (quantifier), 449,10 –> 137,10; Cyanidin-3-O-glucoside (qualifier), 449,10 –> 213,15; ^13^C_9_, ^15^N_1_-Phenylalanine, 176,11 –> 129,25. The Dwell time [ms], collision energy [V], Q1Pre Bias [V] and Q3 Pre Bias [V] were: 32/32/247, –54/-54/-15, –20/-20/-20 and – 14/-22/-20, respectively for Cyanidin-3-O-glucoside (quantifier)/Cyanidin-3-O-glucoside (qualifier)/^13^C_9_, ^15^N_1_-Phenylalanine. The retention time of Cyanidin-3-O-glucoside was 3.38 min and for ^13^C_9_, ^15^N_1_-Phenylalanine 2.52 min. Relative quantification of Cyanidin-3-O-glucoside was done based on the ^13^C_9_, ^15^N_1_-Phenylalanine internal standard from the isotope-labelled amino acid mixture and after normalization to the fresh weight of extracted plant material.

For absolute quantification of Cyanidin-3-O-glucoside and additional analysis of Cyanidin-3-O-(6-O-malonyl-beta-glucoside) the pure sample extracts were further analyzed via HPLC-PDA, which was done on a Shimadzu Nexera XR equipped with a photodiode array detector, a Nucleodur Sphinx RP column (250×4.6 mm, 5µm, Macherey-Nagel) and an EC 4/3 Nucleodur Sphinx RP pre-column (5 µm, Macherey-Nagel). Separation was done in gradient mode with 0.2% formic acid (Fisher Chemical), 0.1% acetonitrile (Fisher Chemical) in water as Solvent A and acetonitrile (Fisher Chemical) as Solvent B, with the column oven set to 20 °C and a flow rate of 1300 µL/min using following solvent settings (Time [min]/B[%]): 0.0/10, 8.0/21, 18.0/49, 18.1/100, 19.0/100, 19.1/10, 24.0/10. Measurement was performed with a PDA detector.

Cyanidin-3-O-glucoside and Cyanidin-3-O-(6-O-malonyl-beta-glucoside) were analyzed at an absorption wavelength of 517 nm and their retention times of 7.055 min and 9.410 min, respectively. Quantification was done based on the comparison with the molar quantity of an external Cyanidin-3-O-glucoside standard curve and after normalization to the fresh weight of extracted plant material.

## Acknowledgments

We thank Ramona Petrig, Pascal Bucher, Lale Bähni, the analytical and teaching laboratory (AuA Lab), Morris Galli, Jesper Wallisch, Max Hofland and Raphael Bossart from Eawag, Marlon Haessler and Benjamin Huessy from the University of Basel, as well as Pascal Poweleit and Alexandra Chávez Argandoña from the University of Mainz for experimental assistance at the pond side and the lab. We thank Yangzi Wang from the University of Mainz for annotating putative gene functions from *D. magna*. Additionally, we thank Holger Schön and David Martín Fernandez from the University of Münster for constructing the swimming boxes.

## Author contributions

Conceptualized the project: CV, DE, SX; Methodology: MS, AM, CW, PS, MS-S, SK, TB, CD-B, ED, LB, CV, DE, SX, JH; Investigation: MS, AM, CW, MS-S, SK, TB, CD-B, ED, JH, LB, CV, DE, SX; Visualization: MS, AM and SX; Supervision CV, DE and SX; Writing—original draft: MS and SX; Writing—review & editing: MS, AM, CV, DE, SX.

## Data and materials availability

Short reads are deposited in NCBI under accession number PRJNA849360. All data and scripts used for generating figures are available on GitHub (https://github.com/Xu-lab-Evolution/EAWAG_ADD/). The raw data are deposited on the Dryad repository (https://datadryad.org/stash/share/Ef3J0yhmOLAlFElcLjh3WZpT9-VRsvahh3b-q4TKa0). Correspondence and requests for materials should be addressed to SX.

## References

1. J. T. Wootton, Predicting direct and indirect effects: An integrated approach using experiments and path analysis. Ecology 75, 151–165 (1994).

2. P. R. Guimaraes, M. M. Pires, P. Jordano, J. Bascompte, J. N. Thompson, Indirect effects drive coevolution in mutualistic networks. Nature 550, 511-+ (2017).

3. L. G. Cosmo et al., Indirect effects shape species fitness in coevolved mutualistic networks. Nature 619, 788–792 (2023).

4. T. M. Knight, M. W. McCoy, J. M. Chase, K. A. McCoy, R. D. Holt, Trophic cascades across ecosystems. Nature 437, 880–883 (2005).

5. A. P. Hendry, A critique for eco-evolutionary dynamics. Funct. Ecol. 33, 84–94 (2019).

6. F. Pelletier, D. Garant, A. P. Hendry, Eco-evolutionary dynamics. Philos T R Soc B 364, 1483-1489 (2009).

7. J. H. Connell, On the prevalence and relative importance of interspecific competition: Evidence from field experiments. Am Nat 122, 661–696 (1983).

8. J. P. Bryant, P. J. Kuropat, S. M. Cooper, K. Frisby, N. Owensmith, Resource availability hypothesis of plant antiherbivore defense tested in a south-african savanna ecosystem. Nature 340, 227–229 (1989).

9. P. D. Coley, J. P. Bryant, F. S. Chapin, 3rd, Resource availability and plant antiherbivore defense. Science 230, 895–899 (1985).

10. R. Karban, A. A. Agrawal, Herbivore offense. Annu Rev Ecol Evol Syst 33, 641–664 (2002).

11. S. Y. Strauss, R. E. Irwin, Ecological and evolutionary consequences of multispecies plant-animal interactions. Annu Rev Ecol Evol Syst 35, 435–466 (2004).

12. D. Lamonica, B. Clement, S. Charles, C. Lopes, Modelling algae-duckweed interaction under chemical pressure within a laboratory microcosm. Ecotox Environ Safe 128, 252–265 (2016).

13. D. Duneau, P. Luijckx, F. Ben-Ami, C. Laforsch, D. Ebert, Resolving the infection process reveals striking differences in the contribution of environment, genetics and phylogeny to host-parasite interactions. BMC Biol. 9, 11 (2011).

14. G. Bento et al., The genetic basis of resistance and matching-allele interactions of a host-parasite system: The *Daphnia magna*-*Pasteuria ramosa* model. PLoS Genet. 13, e1006596 (2017).

15. C. Ameline et al., A two-locus system with strong epistasis underlies rapid parasite-mediated evolution of host resistance. Mol. Biol. Evol. 38, 1512–1528 (2021).

16. W. G. Cochran, Some methods for strengthening the common χ 2 tests. Bioethics 10, 417 (1954).

17. N. Mantel, W. Haenszel, Statistical aspects of the analysis of data from retrospective studies of disease. JNCI: Journal of the National Cancer Institute 22, 719–748 (1959).

18. M. Fredericksen, P. D. Fields, L. Du Pasquier, V. Ricci, D. Ebert, QTL study reveals candidate genes underlying host resistance in a Red Queen model system. PLoS Genet. 19, e1010570 (2023).

19. J.-Y. Choi et al., Population growth of the cladoceran, *Daphnia magna*: a quantitative analysis of the effects of different algal food. PLoS One 9, e95591 (2014).

20. D. S. Glazier, Effects of food, genotype, and maternal size and age on offspring investment in *Daphnia magna*. Ecology 73, 910–926 (1992).

21. K. N. Hoefnagel, E. de Vries, E. Jongejans, W. Verberk, The temperature-size rule in *Daphnia magna* across different genetic lines and ontogenetic stages: Multiple patterns and mechanisms. Ecol Evol 8, 3828–3841 (2018).

22. L. B. Goss, D. L. Bunting, Daphnia development and reproduction: Responses to temperature. J. Therm. Biol. 8, 375–380 (1983).

23. C. Ameline et al., Genetic slippage after sex maintains diversity for parasite resistance in a natural host population. Sci Adv 8, eabn0051 (2022).

24. A. Narwani et al., Interactive effects of foundation species on ecosystem functioning and stability in response to disturbance. P Roy Soc B-Biol Sci 286, 20191857 (2019).

25. K. J. Appenroth, G. Nickel, Turion formation in *Spirodela polyrhiza*: The environmental signals that induce the developmental process in nature. Physiol Plantarum 138, 312–320 (2010).

26. L. Böttner et al., Herbivory can increase plant fitness via reduced interspecific competition – evidence from models and mesocosms. bioRxiv, 2024.2004.2020.589392 (2024).

27. P. Angst et al., Genetic drift shapes the evolution of a highly dynamic metapopulation. Mol. Biol. Evol. 39, (2022).

28. H. Utermöhl, Neue Wege in der quantitativen Erfassung des Plankton.(Mit besonderer Berücksichtigung des Ultraplanktons.). SIL Proceedings 5, 567–596 (1931).

29. D. Duneau et al., Stochastic variation in the initial phase of bacterial infection predicts the probability of survival in *D. melanogaster*. eLife 6, (2017).

30. Felix Krueger et al. (Zenodo, 2023), vol. 2024.

31. L. Cornetti, P. D. Fields, L. Du Pasquier, D. Ebert, Long-term balancing selection for pathogen resistance maintains trans-species polymorphisms in a planktonic crustacean. Nat Commun 15, 5333 (2024).

32. H. Li, R. Durbin, Fast and accurate short read alignment with Burrows-Wheeler transform. Bioinformatics 25, 1754–1760 (2009).

33. P. Danecek et al., Twelve years of SAMtools and BCFtools. Gigascience 10, (2021).

34. A. Iranmehr, A. Akbari, C. Schlotterer, V. Bafna, Clear: Composition of likelihoods for evolve and resequence experiments. Genetics 206, 1011-1023 (2017).

35. R. Kofler, R. V. Pandey, C. Schlotterer, PoPoolation2: identifying differentiation between populations using sequencing of pooled DNA samples (Pool-Seq). Bioinformatics 27, 3435–3436 (2011).

36. M. Gautier, R. Vitalis, L. Flori, A. Estoup, f-Statistics estimation and admixture graph construction with Pool-Seq or allele count data using the R package poolfstat. Mol Ecol Resour 22, 1394–1416 (2022).

37. K. Luu, E. Bazin, M. G. Blum, pcadapt: an R package to perform genome scans for selection based on principal component analysis. Mol Ecol Resour 17, 67–77 (2017).

38. N. Mantel, W. Haenszel, Statistical aspects of the analysis of data from retrospective studies of disease. J. Natl. Cancer Inst. 22, 719–748 (1959).

39. W. G. Cochran, Some methods for strengthening the common χ² tests. J Am Stat Assoc 50, 572–572 (1955).

40. M. E. Brooks et al., glmmTMB balances speed and flexibility among packages for zero-inflated generalized linear mixed modeling. R J 9, 378–400 (2017).

41. D. E. Wood, J. Lu, B. Langmead, Improved metagenomic analysis with Kraken 2. Genome Biol 20, (2019).

42. A. Thivolle, M. Paljakka, D. Ebert, P. D. Fields, The genome of Pasteuria ramosa reveals a high turnover rate of collagen-like genes. bioRxiv, 2024.2002.2009.579640 (2024).

43. A. Malacrino et al., Induced responses contribute to rapid adaptation of *Spirodela polyrhiza* to herbivory by *Lymnaea stagnalis*. Commun Biol 7, 81 (2024).

