## Supplementary information for "Terrestrial herbivory drives adaptive evolution in an aquatic community via indirect effects"

**Supplementary figures and tables**

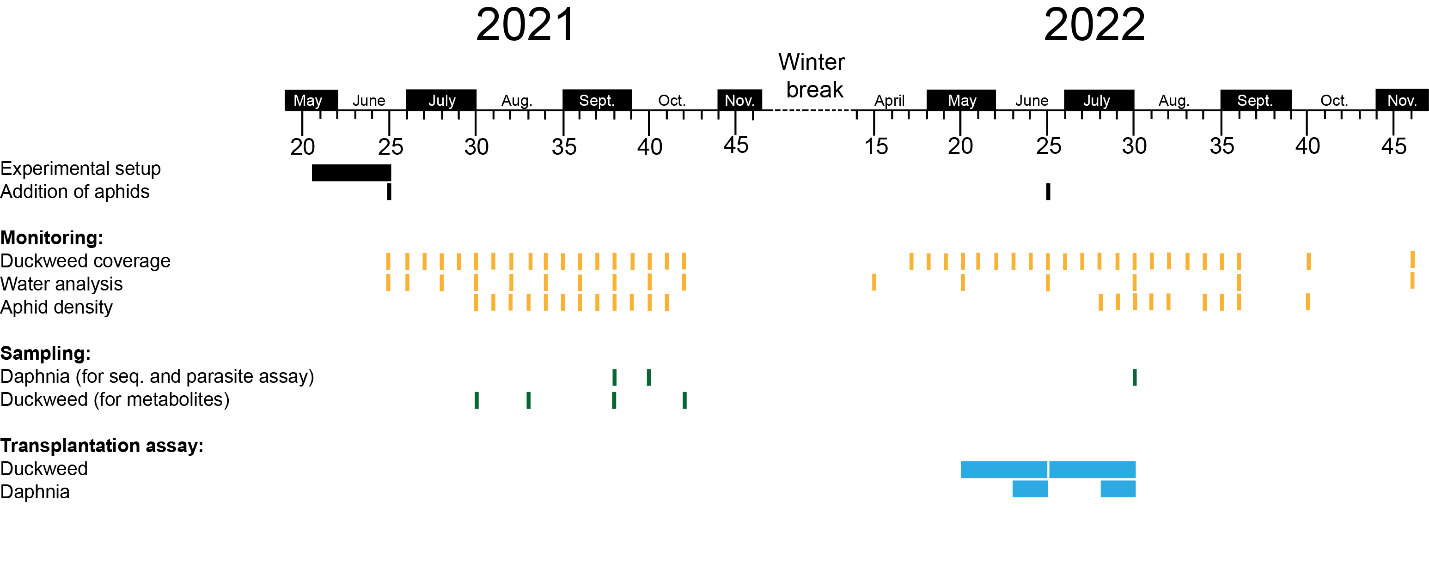

**Fig. S1 Chronological outline of the experimental setup and sampling.** Month and calendar week information is provided above and below the timeline, respectively.

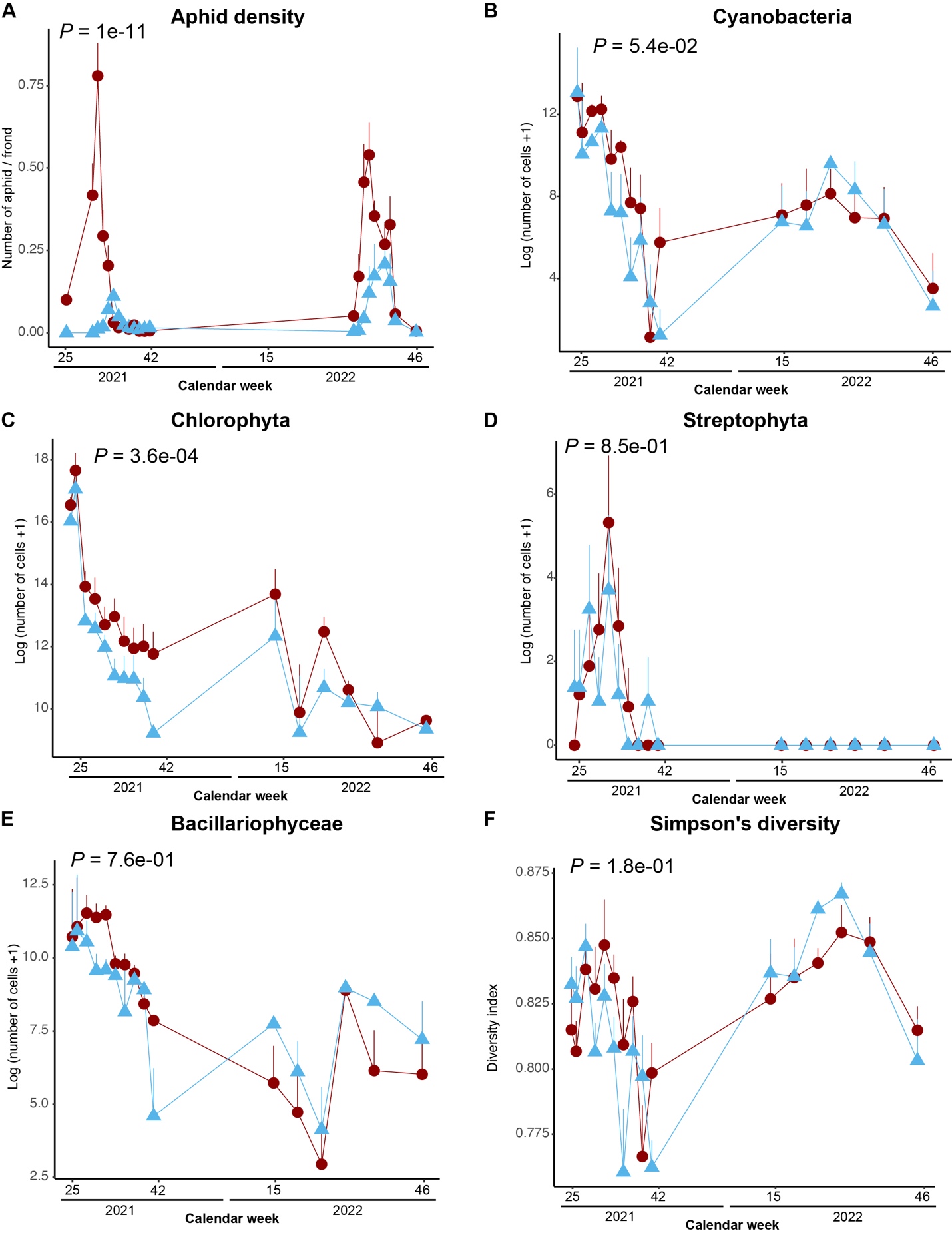

**Fig. S2 Population changes among different species in the community.** A-E refer to the abundance of aphid (A), Cyanobacteria (B), Chlorophyta (C), Streptophyta (D) and Bacillariophyceae (E). Panel F refers to phytoplankton diversity in the aquatic community, which is calculated using the log-transformed abundance (number of cells per liter) of each phytoplankton group. P-values refer to the effects of aphid herbivory. All P-values were estimated using mixed-effects models with time and pond block as random factors. Light blue and red colors refer to control and aphid herbivory ponds. Error bar refers to standard errors.

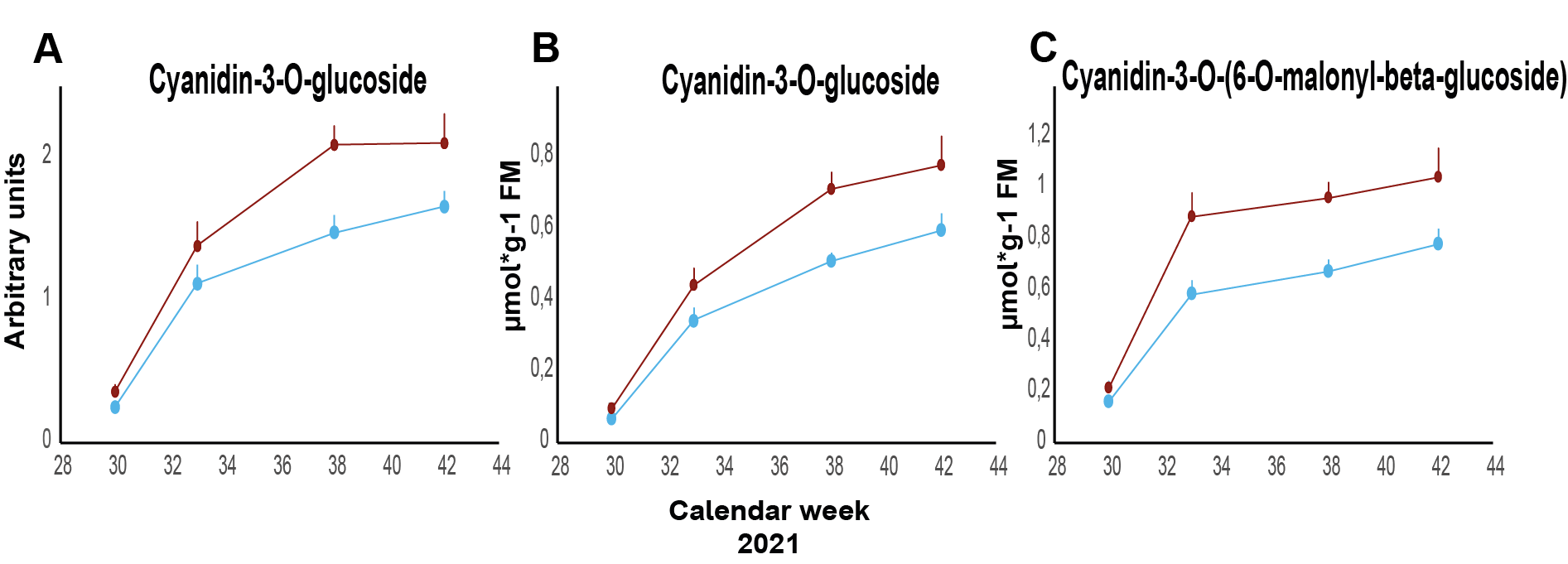

**Fig. S3 Anthocyanin abundance in the duckweed populations.**

The abundance of Cyanidin-3-O-glucoside (A, B) and Cyanidin-3-O-(6-O-malonyl-beta-glucoside) (C) in the duckweed plants at different time points in 2021. Light blue and red colors refer to control and aphid herbivory ponds. Data were either measured by LC-MS (A; relative quantification) or HPLC-PDA (B, C; absolute quantification). Error bars refer to standard errors.

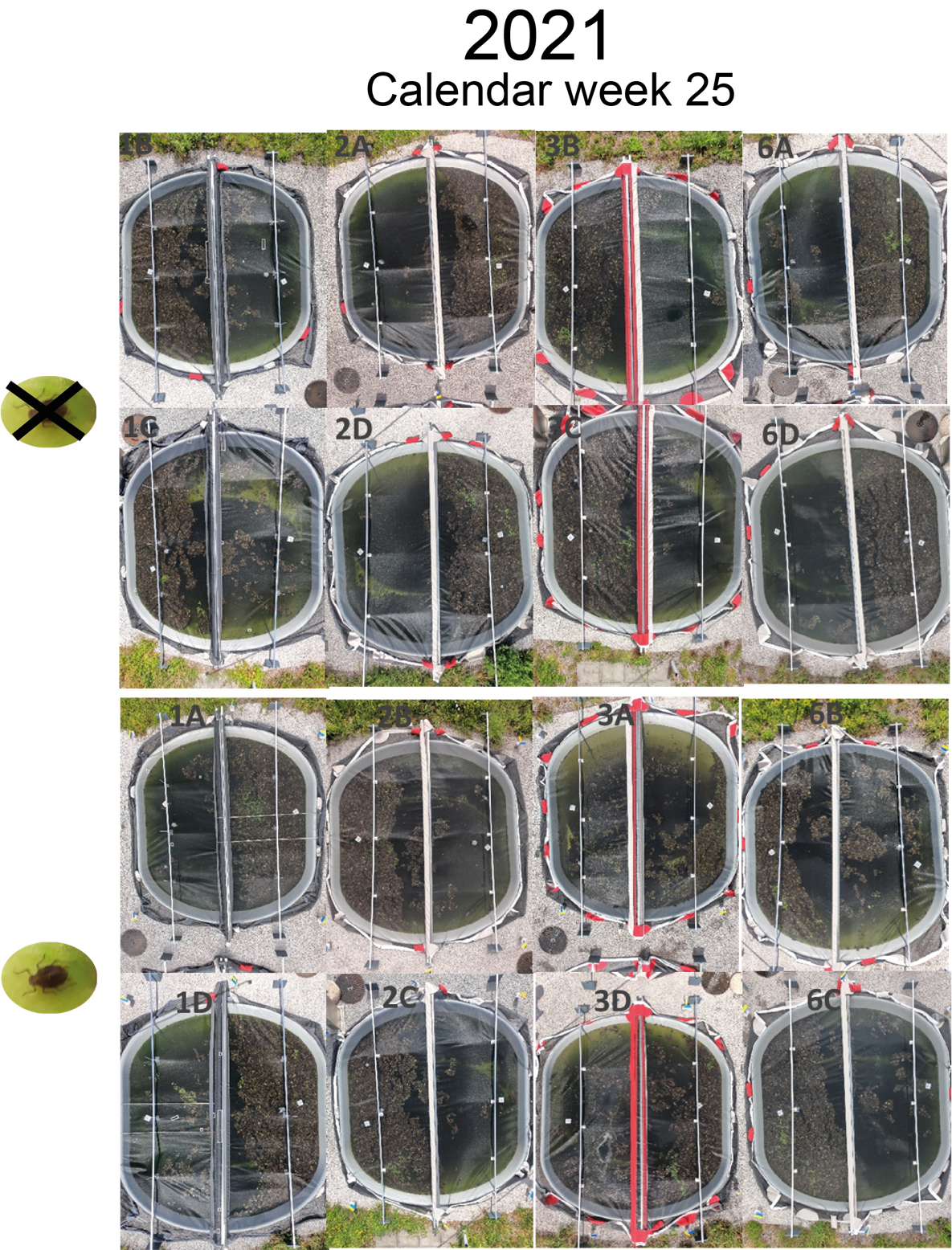

**Fig. S4. Overview picture of the ponds at calendar week 25 in 2021.** The pond ID is indicated in gray. In the upper part, the control ponds are shown, and in the lower part, the aphid herbivory ponds.

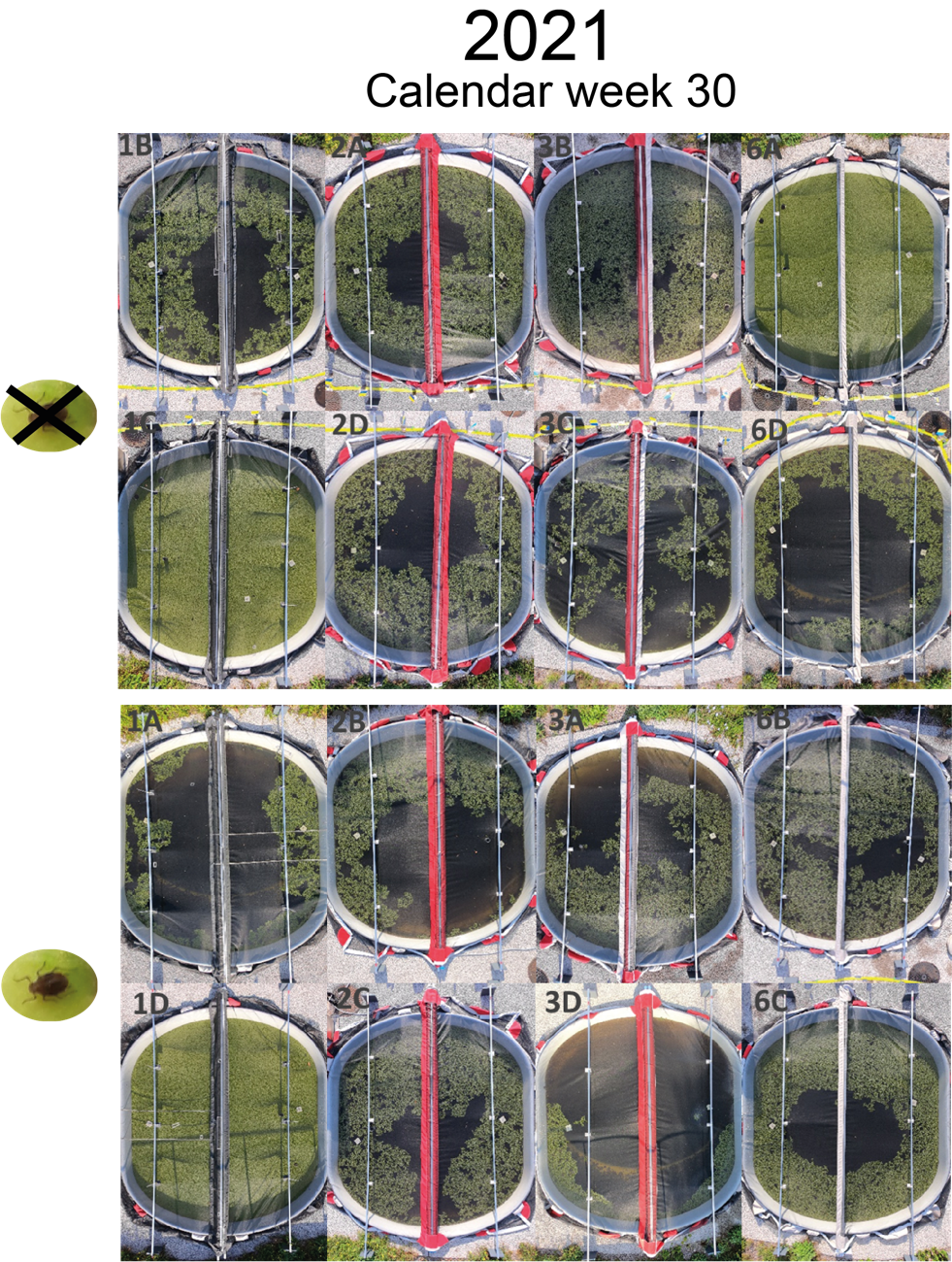

**Fig. S5 Overview picture of the ponds at calendar week 30 in 2021** The pond ID is indicated in gray. In the upper part, the control ponds are shown, and in the lower part, the aphid herbivory ponds.

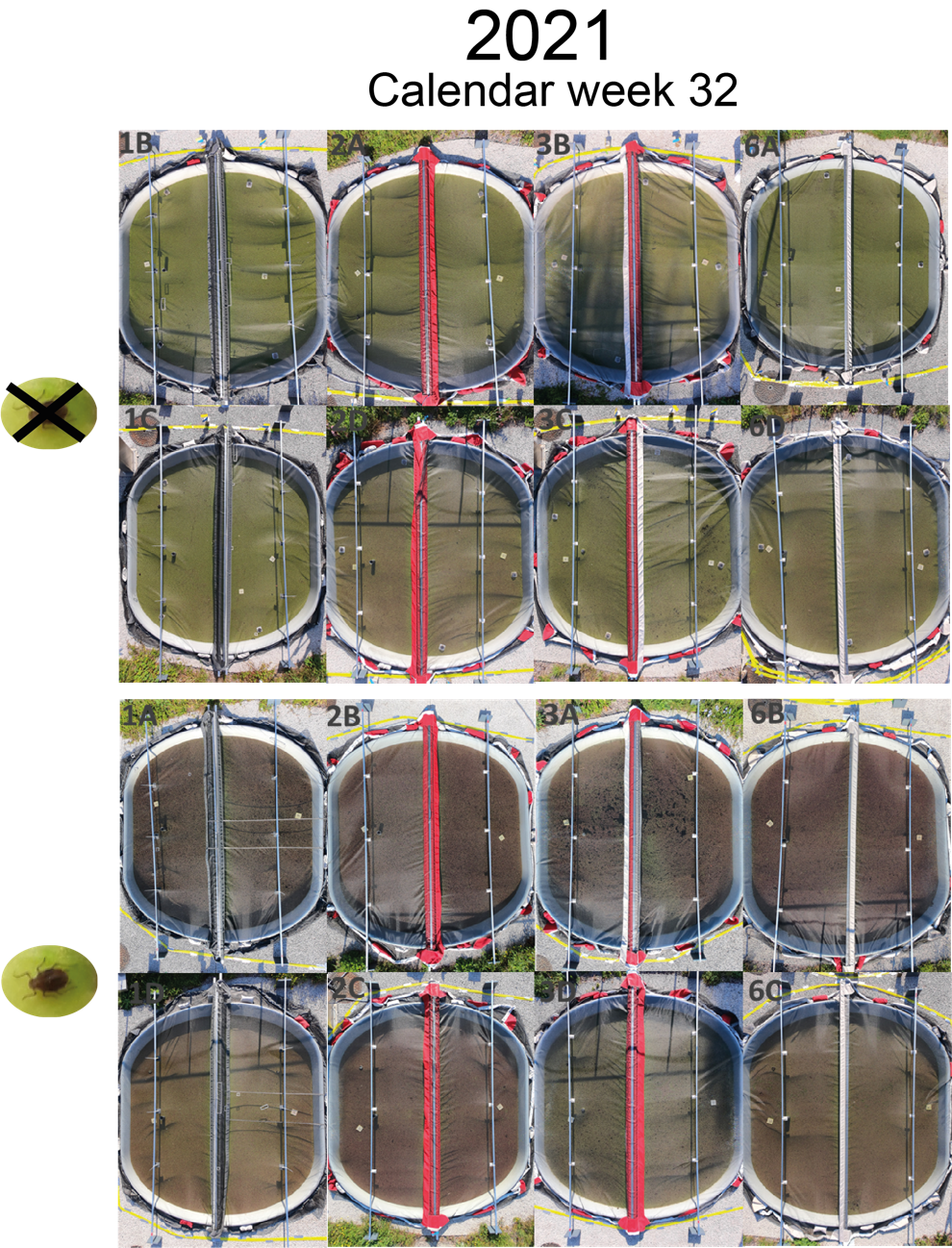

**Fig. S6 Overview picture of the ponds at calendar week 32 in 2021** The pond ID is indicated in gray. In the upper part, the control ponds are shown, and in the lower part, the aphid herbivory ponds.

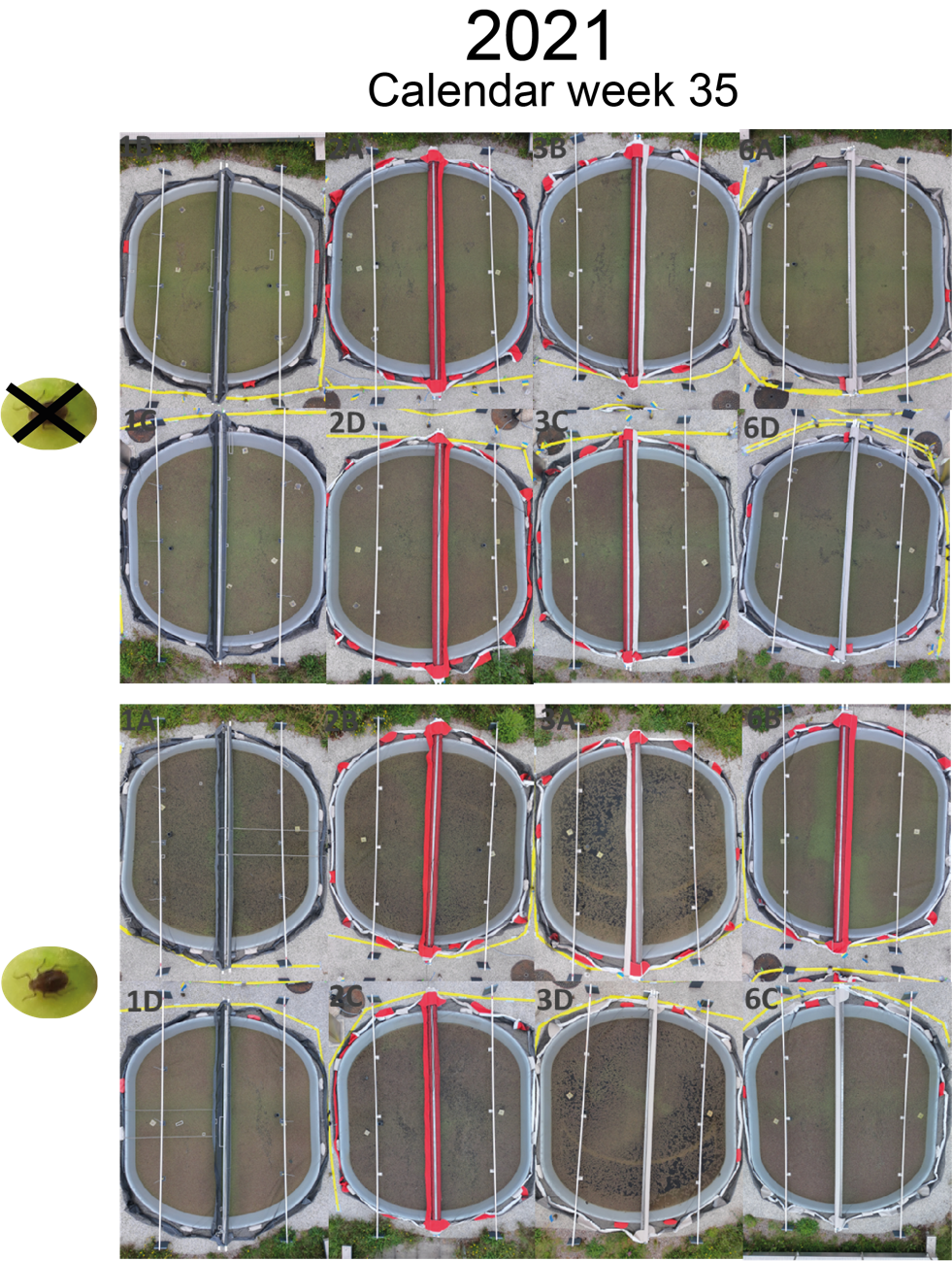

**Fig. S7 Overview picture of the ponds at calendar week 35 in 2021** The pond ID is indicated in gray. In the upper part, the control ponds are shown, and in the lower part, the aphid herbivory ponds.

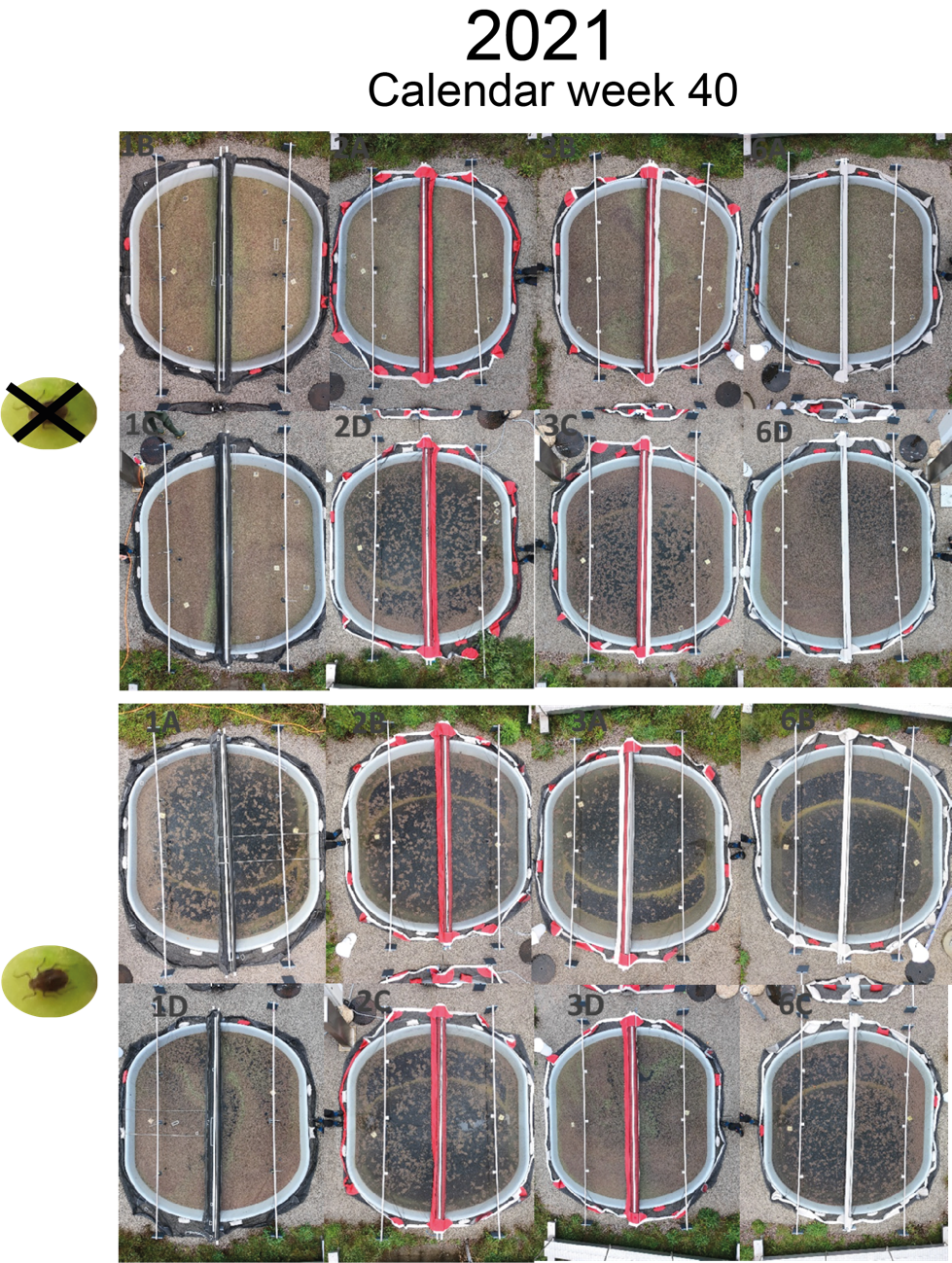

**Fig. S8 Overview picture of the ponds at calendar week 40 in 2021** The pond ID is indicated in gray. In the upper part, the control ponds are shown, and in the lower part, the aphid herbivory ponds.

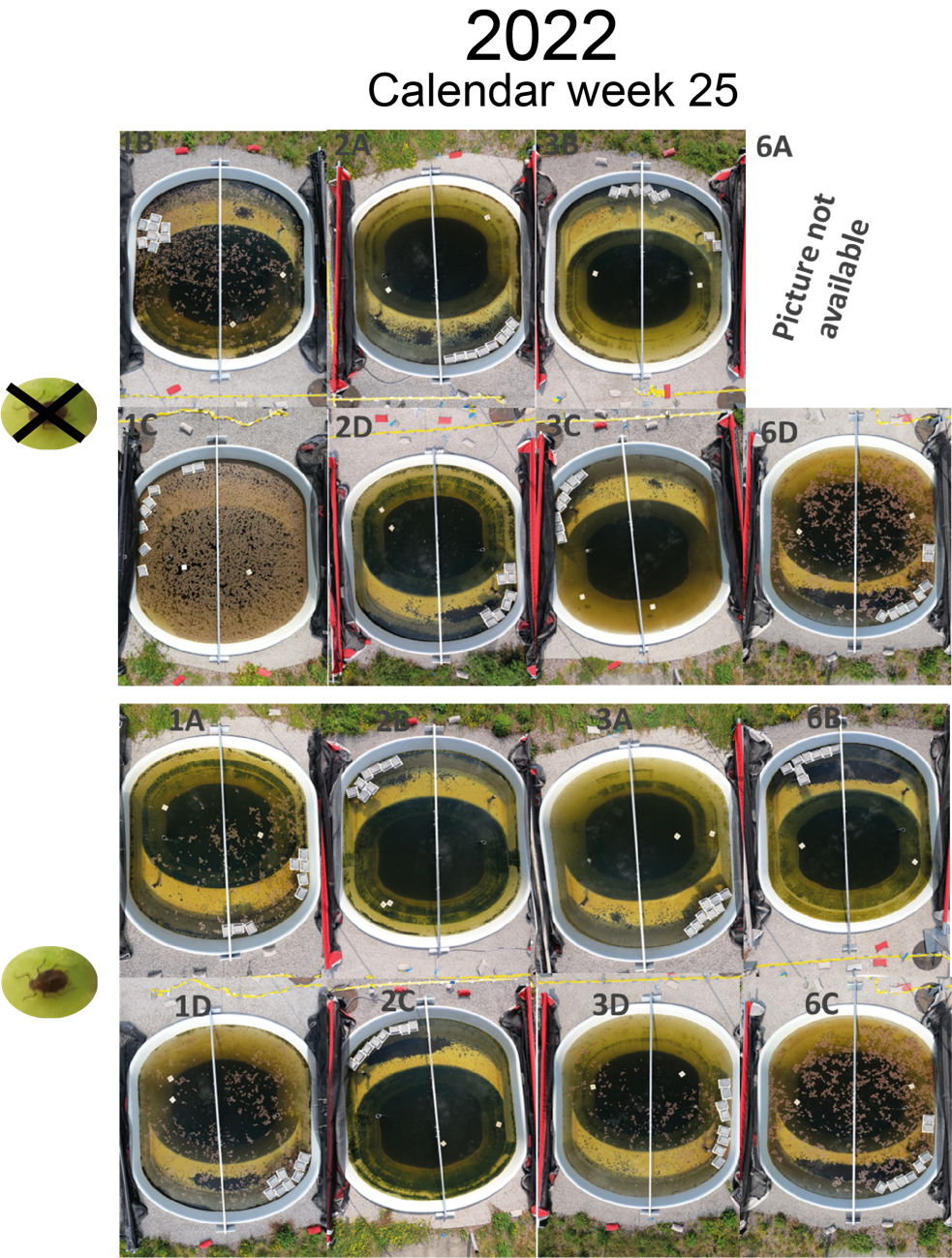

**Fig. S9 Overview picture of the ponds at calendar week 25 in 2022** The pond ID is indicated in gray. In the upper part, the control ponds are shown, and in the lower part, the aphid herbivory ponds.

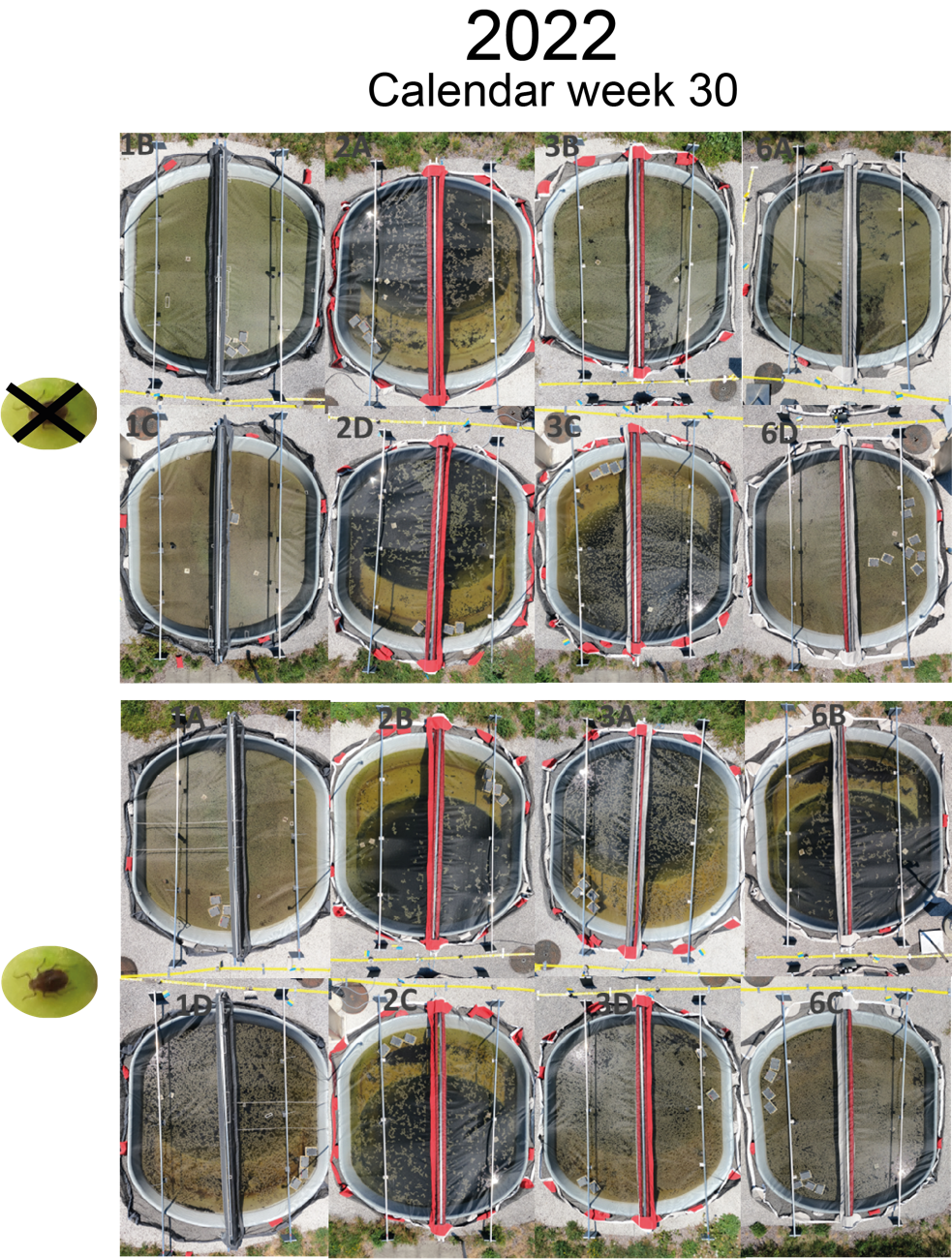

**Fig. S10 Overview picture of the ponds at calendar week 30 in 2022** The pond ID is indicated in gray. In the upper part, the control ponds are shown, and in the lower part, the aphid herbivory ponds.

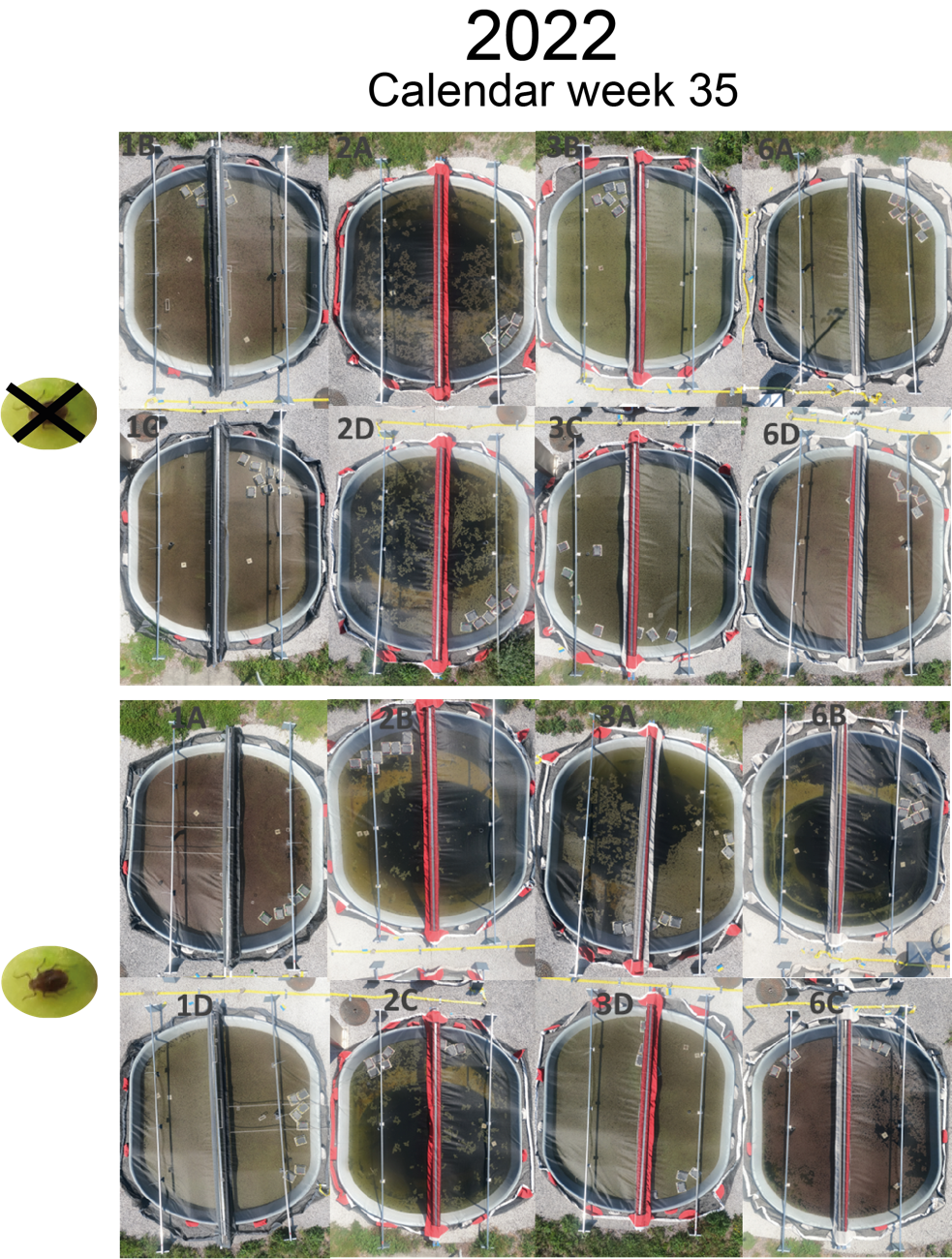

**Fig. S11 Overview picture of the ponds at calendar week 35 in 2022** The pond ID is indicated in gray. In the upper part, the control ponds are shown, and in the lower part, the aphid herbivory ponds.

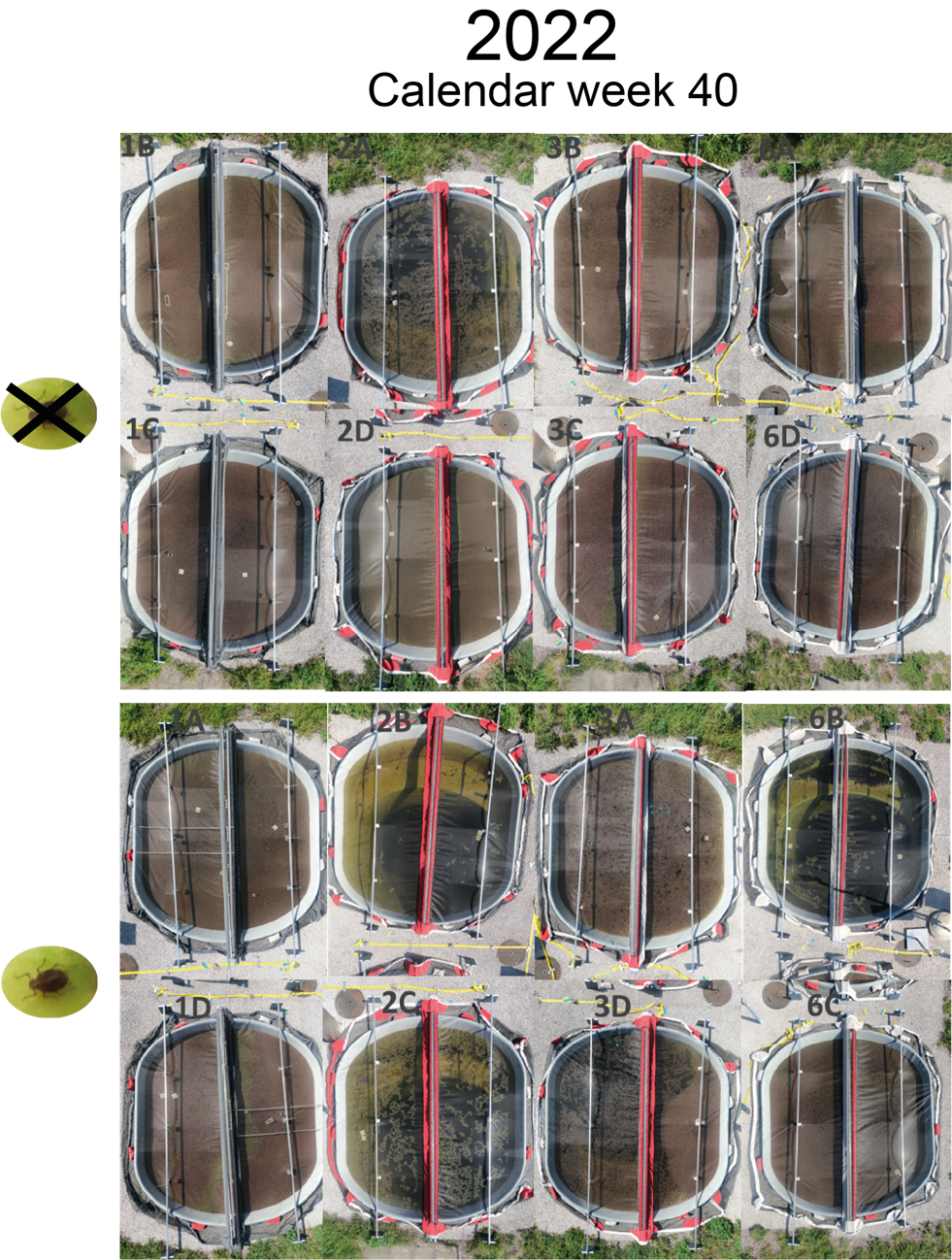

**Fig. S12 Overview picture of the ponds at calendar week 40 in 2022** The pond ID is indicated in gray. In the upper part, the control ponds are shown, and in the lower part, the aphid herbivory ponds.

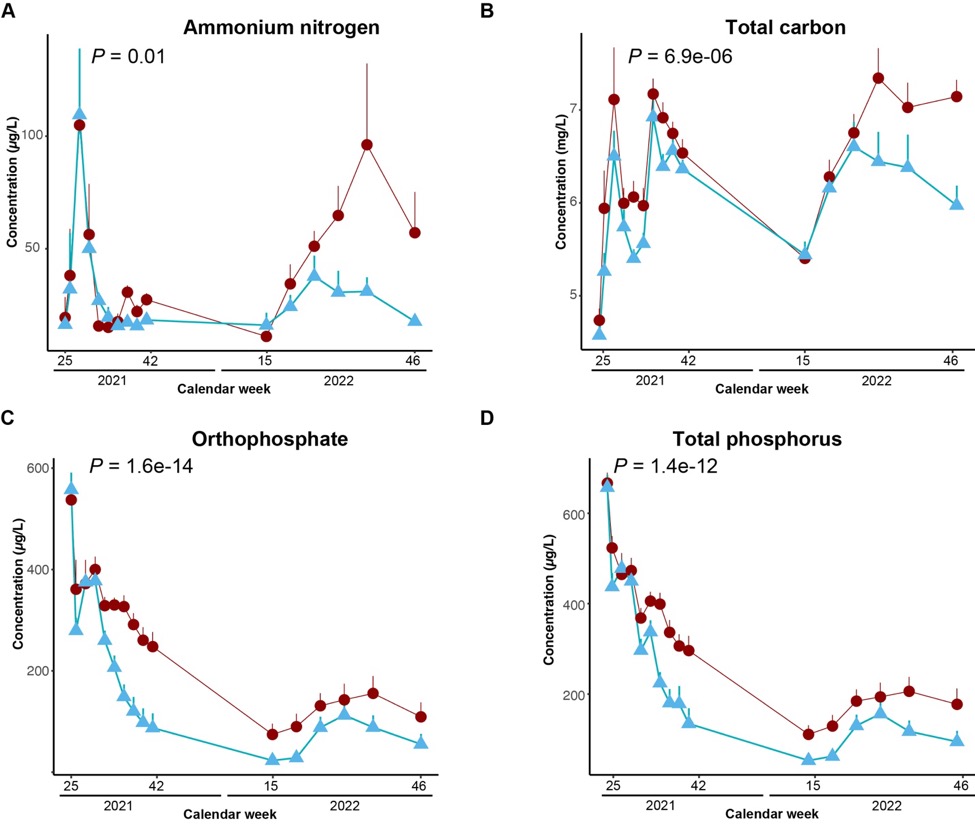
 **Fig. S13 Nutrient levels in the aquatic environment.** A-D refer to the levels of ammonium nitrogen (A), total carbon (B), orthophosphate (C), and total phosphorus (D) in the water. P-values refer to the effects of aphid herbivory. All P-values were estimated using mixed-effects models with time and pond block as random factors. Light blue and red colors refer to control and aphid herbivory ponds. Error bar refers to standard errors.

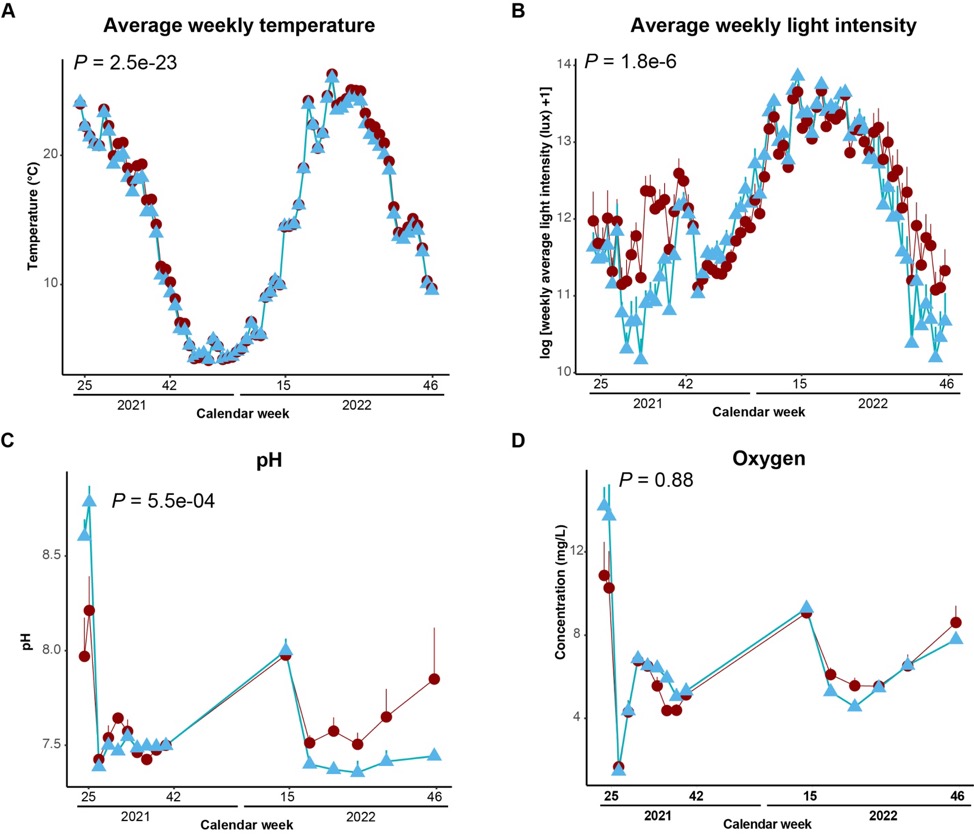

**Fig. S14 Temperature, light intensity, pH and oxygen levels in the ponds.** A-D refer to the average weekly temperature (A), weekly average of the daily light intensity (B), pH (C), and dissolved oxygen levels (D). P-values refer to the effects of aphid herbivory. All P-values were estimated using mixed-effects models with time and pond block as random factors. Light blue and red colors refer to control and aphid herbivory ponds. Error bars refer to standard errors.

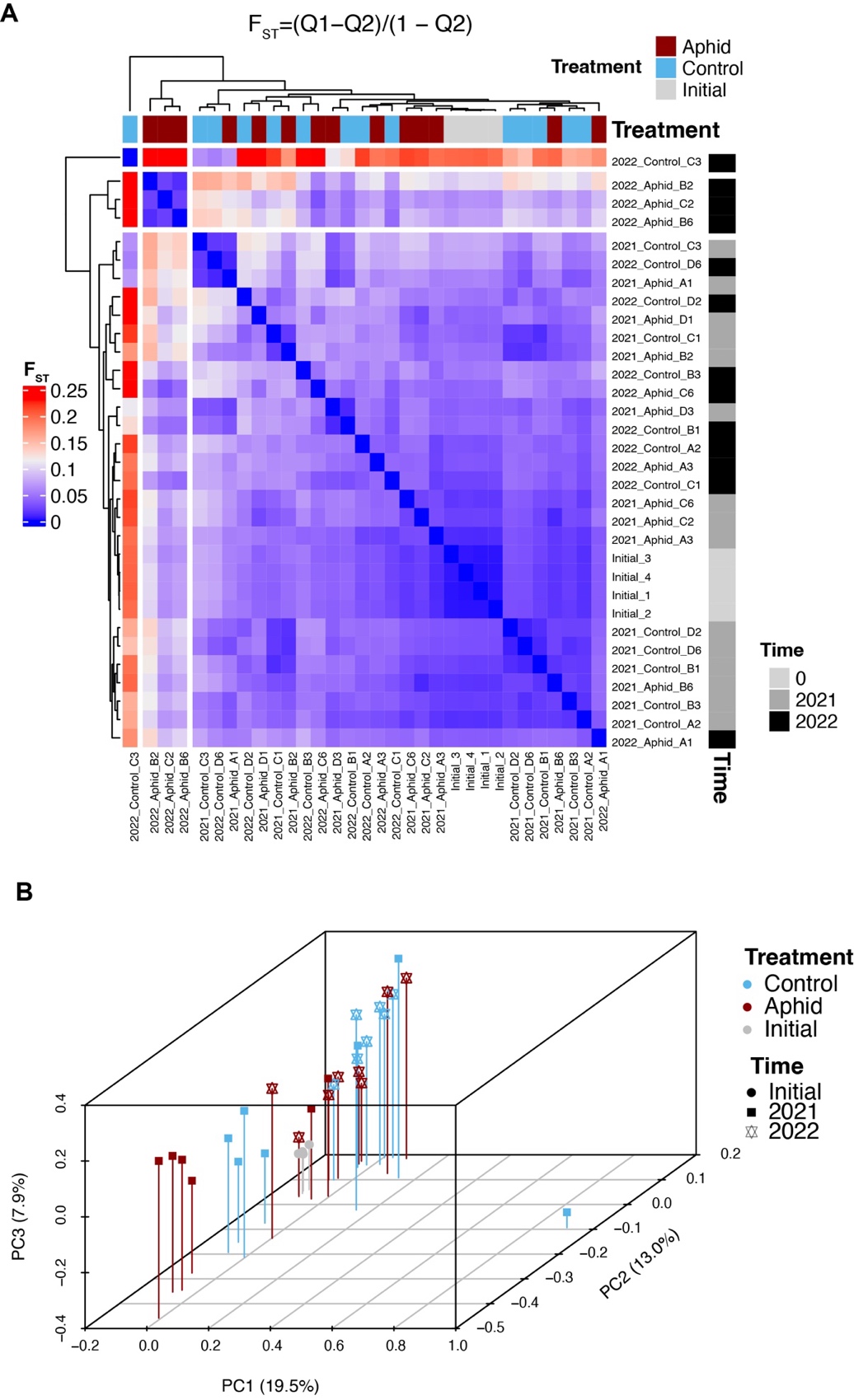

**Fig. S15 Genomic differentiation among populations.** A: heatmap shows the similarity among *D. magna* populations based on pairwise F_st_. B: principle component plot showing the variations among evolved and initial *D. magna* populations. Among all samples, pond 3C in 2022 showed the most difference from the others. The four initial populations showed the lowest F_st_ among each other. Populations from 2022 showed increased F_st_ to other populations.

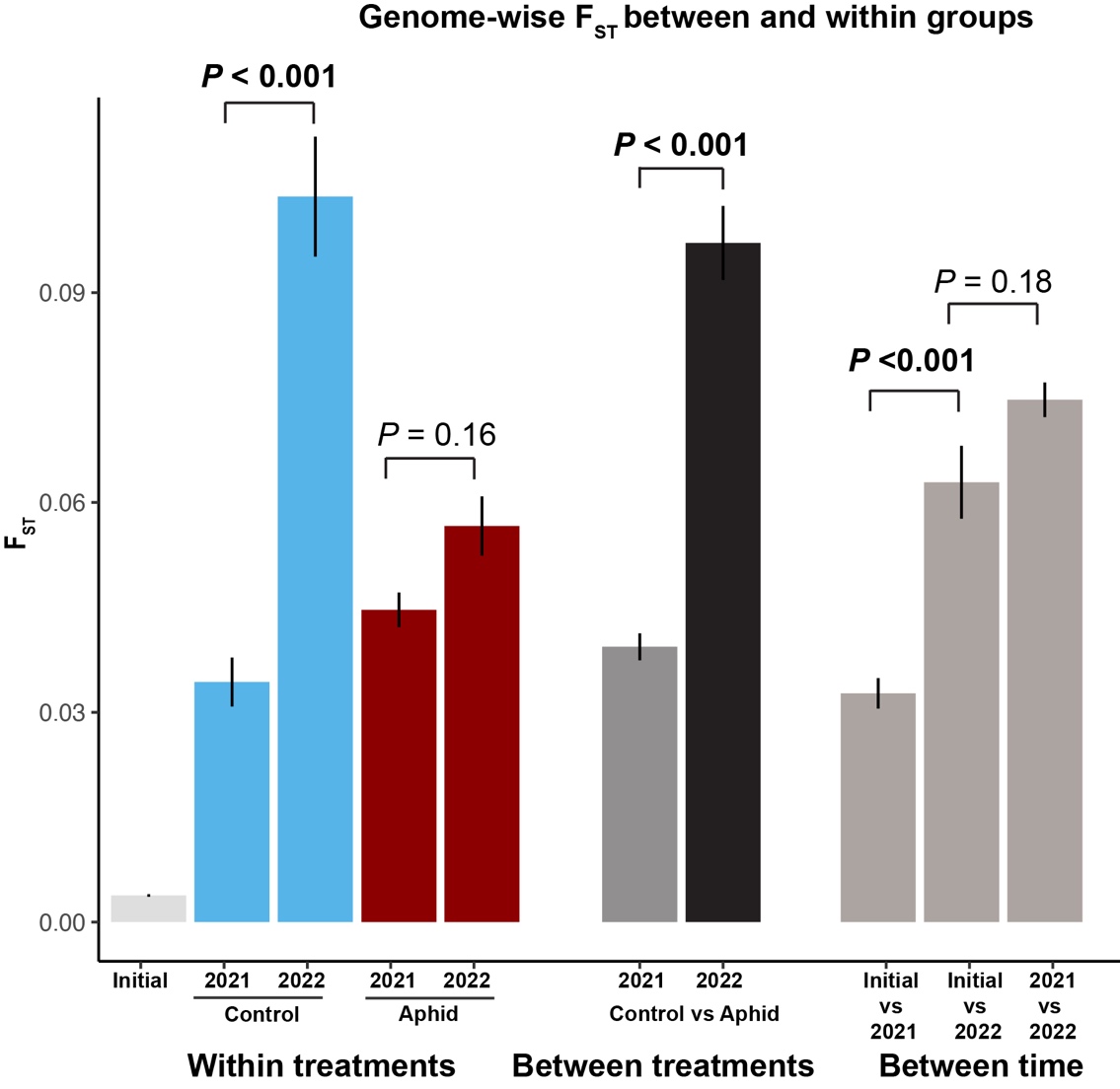

**Fig. S16 Genomic differentiation of *D. magna* within and between treatments and time.** The y-axis refers to the Fst values. The x-axis refers to different comparisons. Mean and standard errors are shown. P-values were estimated using one-way ANOVA.

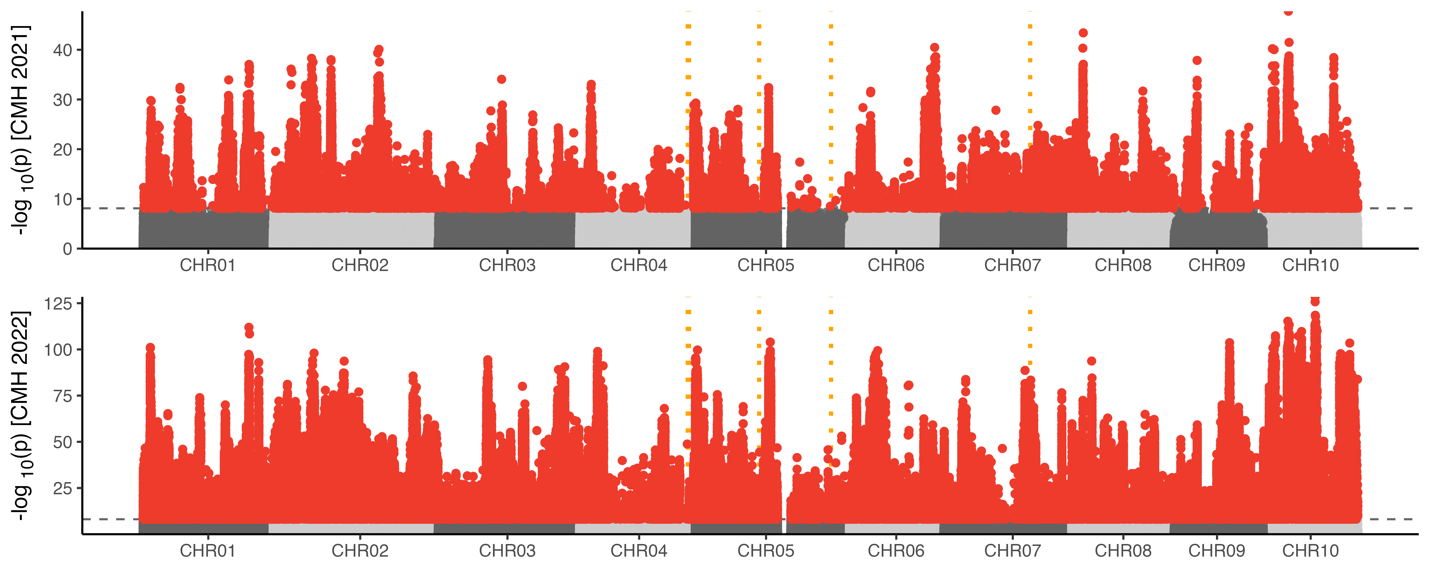

**Fig. S17 Cochran–Mantel–Haenszel test showing *D. magna* genomic divergence between control and aphid-herbivory populations.** The y-axis refers to -log_10_ P-values. The x-axis refers to the genomic position of each SNP. The horizontal dashed lines show the *P* < 0.05 cutoffs after Bonferroni correction. Significant SNPs are shown in red. The vertical lines in orange refer to the location of previously identified loci that affect *Pasteuria ramosa* attachment ability to *D. magna*. Samples collected in 2021 and 2022 are shown in upper and lower panels, respectively.

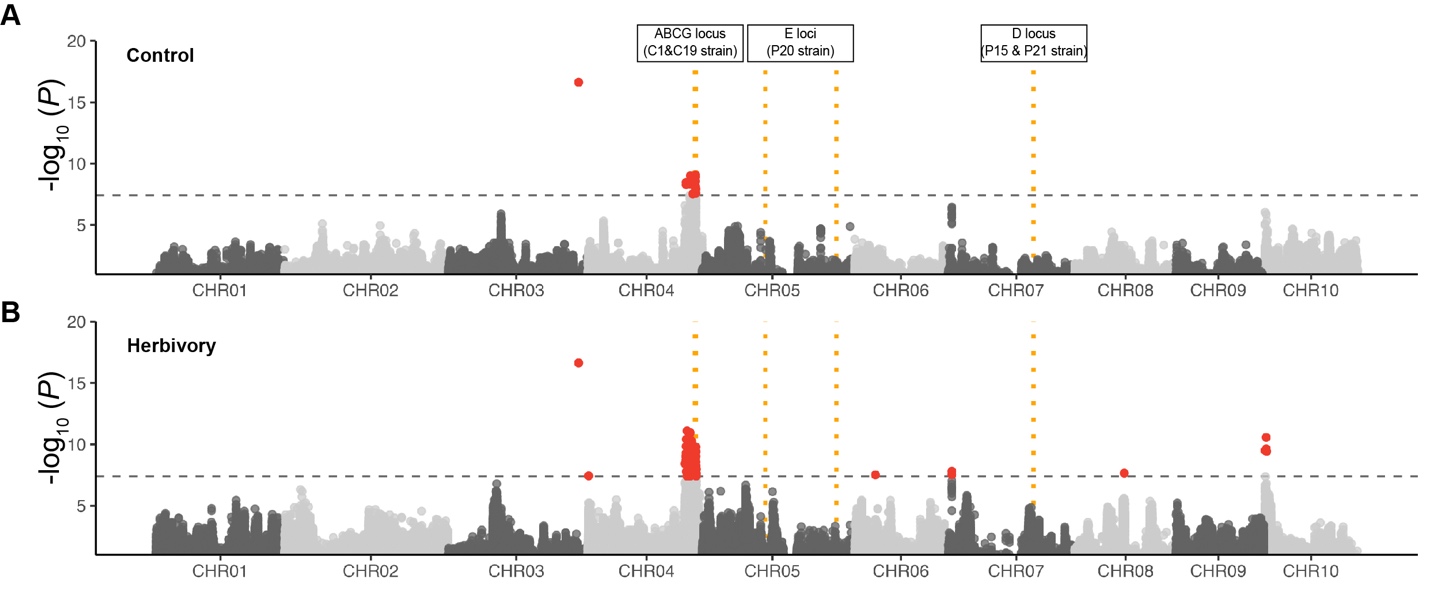

**Fig. S18 Genomic selection on the *D. magna* populations.** The y-axis refers to -log_10_ P-values. X-axis refers to the genomic position of each SNP. The populations evolved in control (A) and aphid herbivory ponds (B) are shown in the upper and lower panels, respectively. The horizontal dashed lines show the *P* < 0.05 cutoff after Bonferroni correction. Significant SNPs are shown in red. The vertical lines in orange refer to the location of previously identified loci that affect *Pasteuria ramosa* attachment ability to *D. magna*.

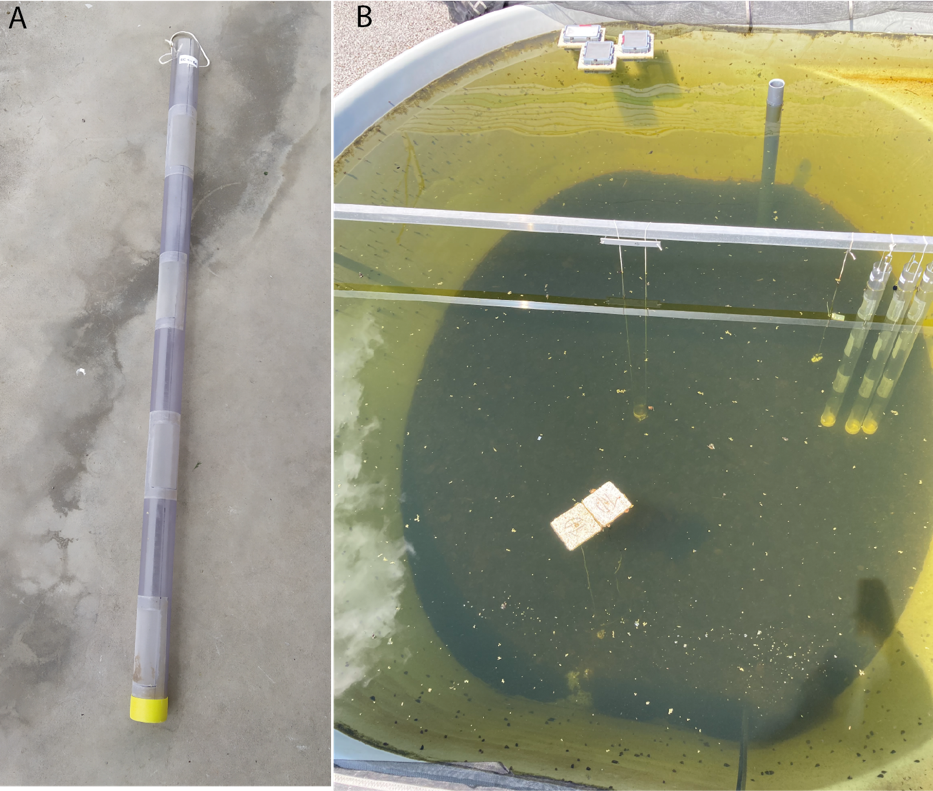

**Fig. S19 Setup of the *Daphnia magna* transplant experiments.** (A) For the *D. magna* transplant experiment, we used PVC columns that contained various cut-outs covered by mesh to allow for an exchange of water and phytoplankton while retaining *D. magna*. (B) PVC columns containing the *Daphnia* were placed in the ponds next to each other by attaching them to the metal bar in the middle of the ponds.

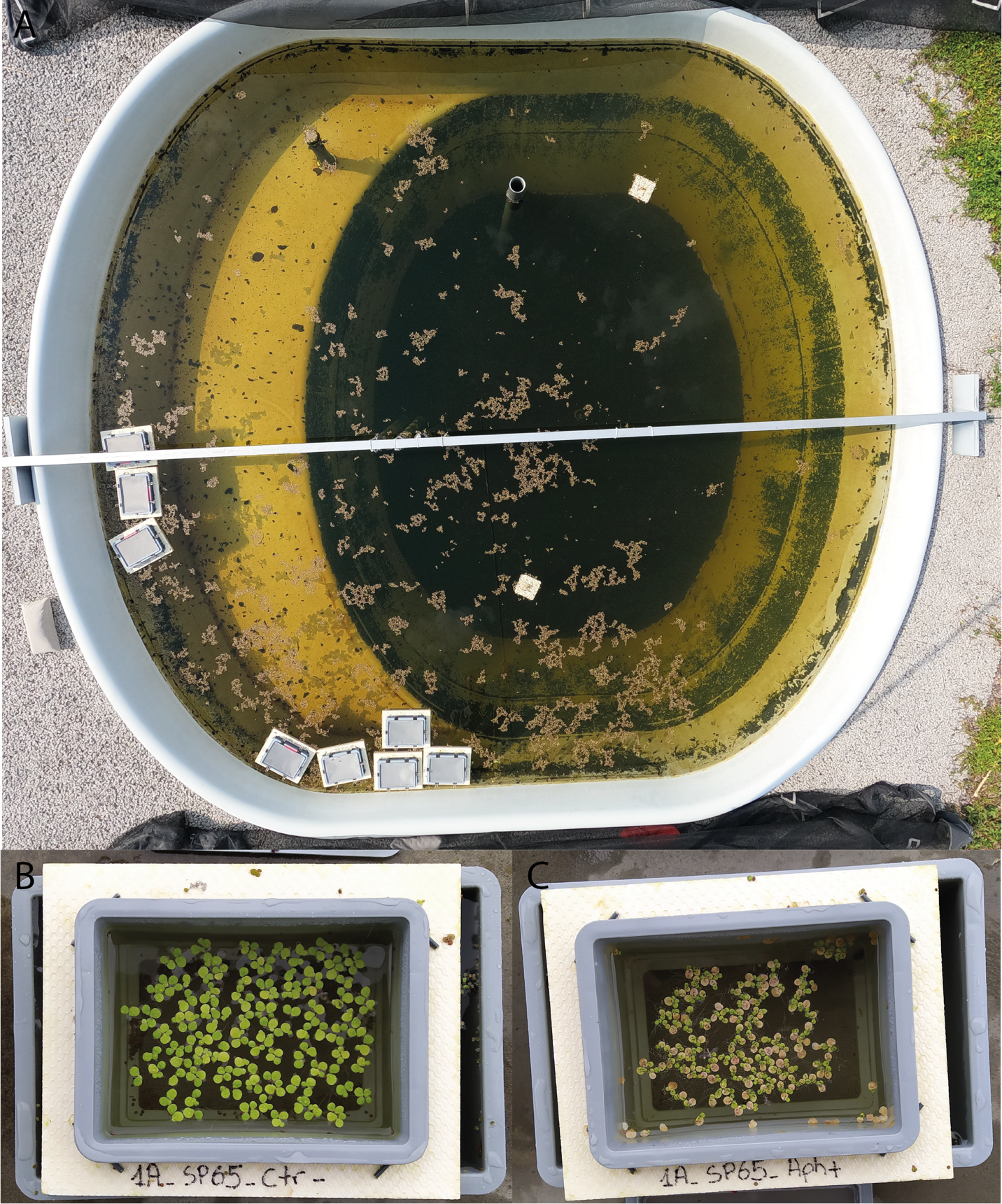

**Fig. S20 Setup of the *Spirodela polyrhiza* and aphid transplant assays within the ponds.** (A) Swimming boxes containing the duckweed of the transplant experiment were allowed to move freely on the water. Exemplary swimming boxes with control (B) and aphid herbivory treatment (C). During sampling and analysis, boxes were placed in a slightly bigger, closed box filled with pond water to temporarily take them out of the ponds (B, C).

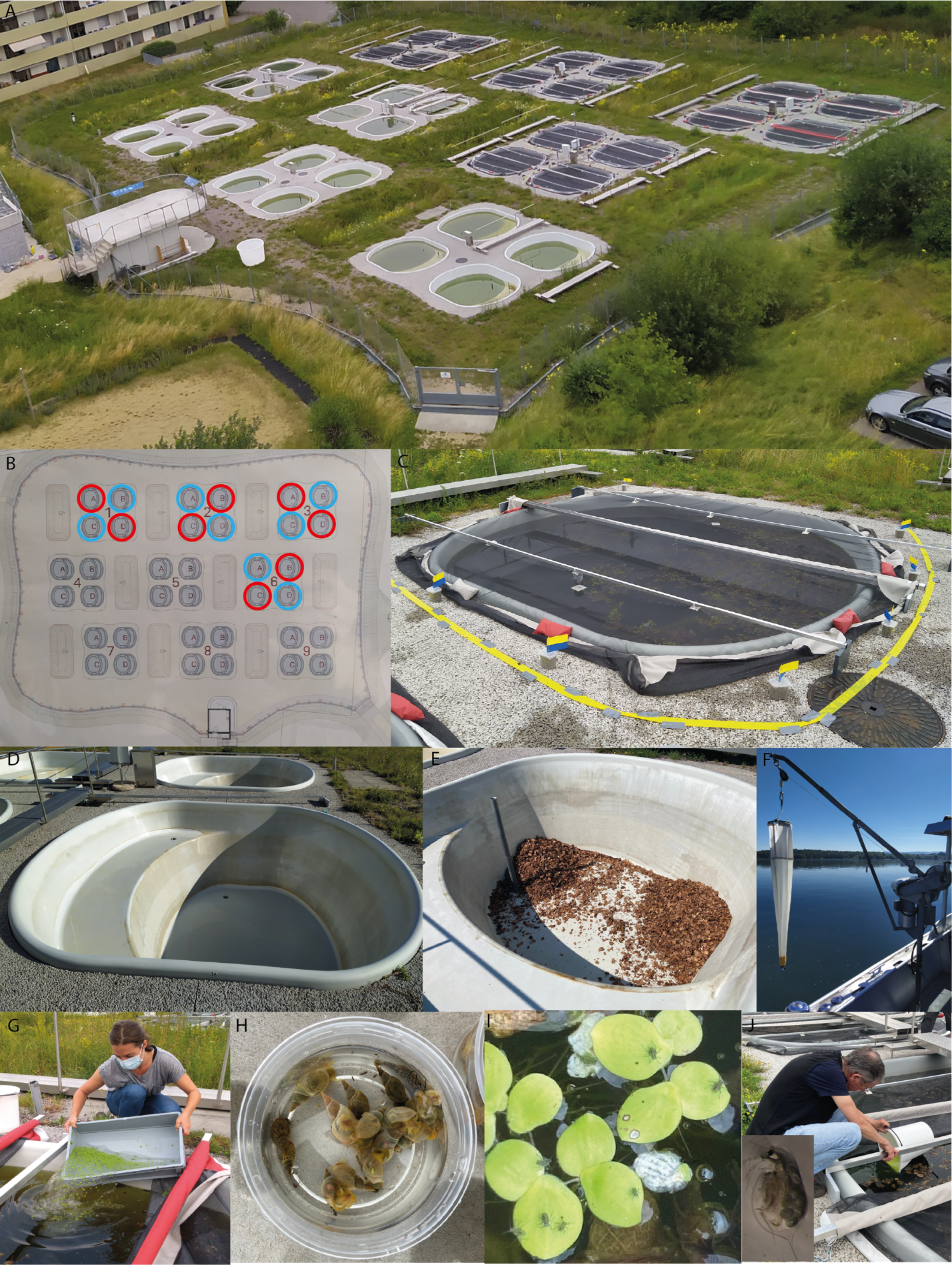

**Fig. S21 Experimental setup**. (A) Overview of the pond facility at Eawag. (B) Scheme of the used ponds and their designation. Ponds used for the experiment are highlighted. Light blue and red colors refer to control and aphid herbivory ponds. (C) Close-up of a fully prepared pond with mesh cover and the aphid protection measures (yellow and blue sticky traps) around. (D) Empty pond before. (E) Pond with leaf litter. (F) Equipment used to sample the plankton within water columns at Greifensee for the inoculation of the ponds at the beginning of the experiment. Addition of (G) duckweed, (H) snails, (I) aphids, and (J) daphnia at the start of the experiment.

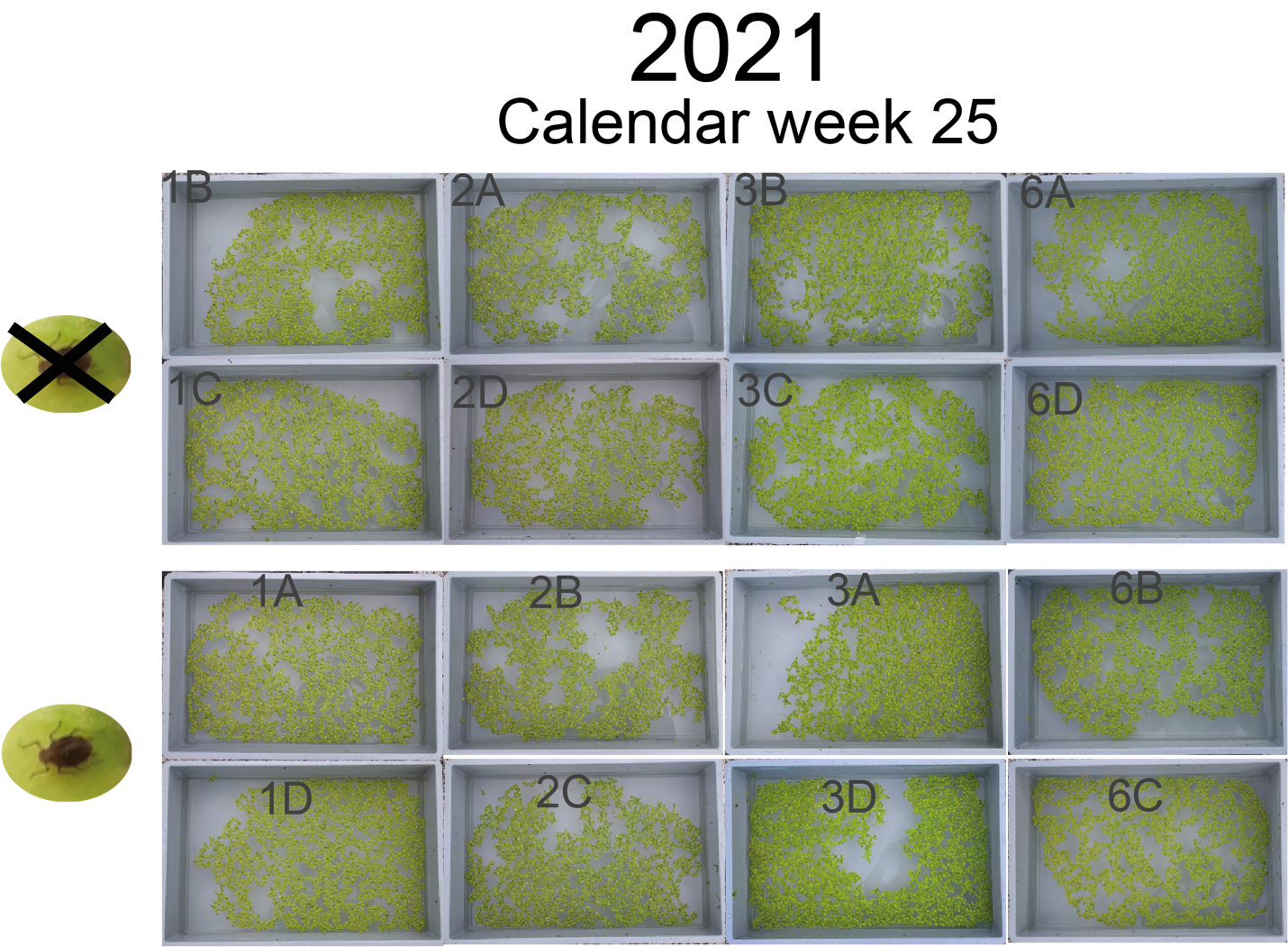

**Fig. S22 Overview picture of the duckweed used for the pond inoculation at the start of the experiment.** *Spirodela polyrhiza* fronds before addition to the ponds in calendar week 25 in 2021. The pond ID which the material corresponds to is indicated in gray. In the upper part, the material for the control ponds is shown, and in the lower part, those of the aphid herbivory ponds. Pond 6A was removed from the analysis in 2021.

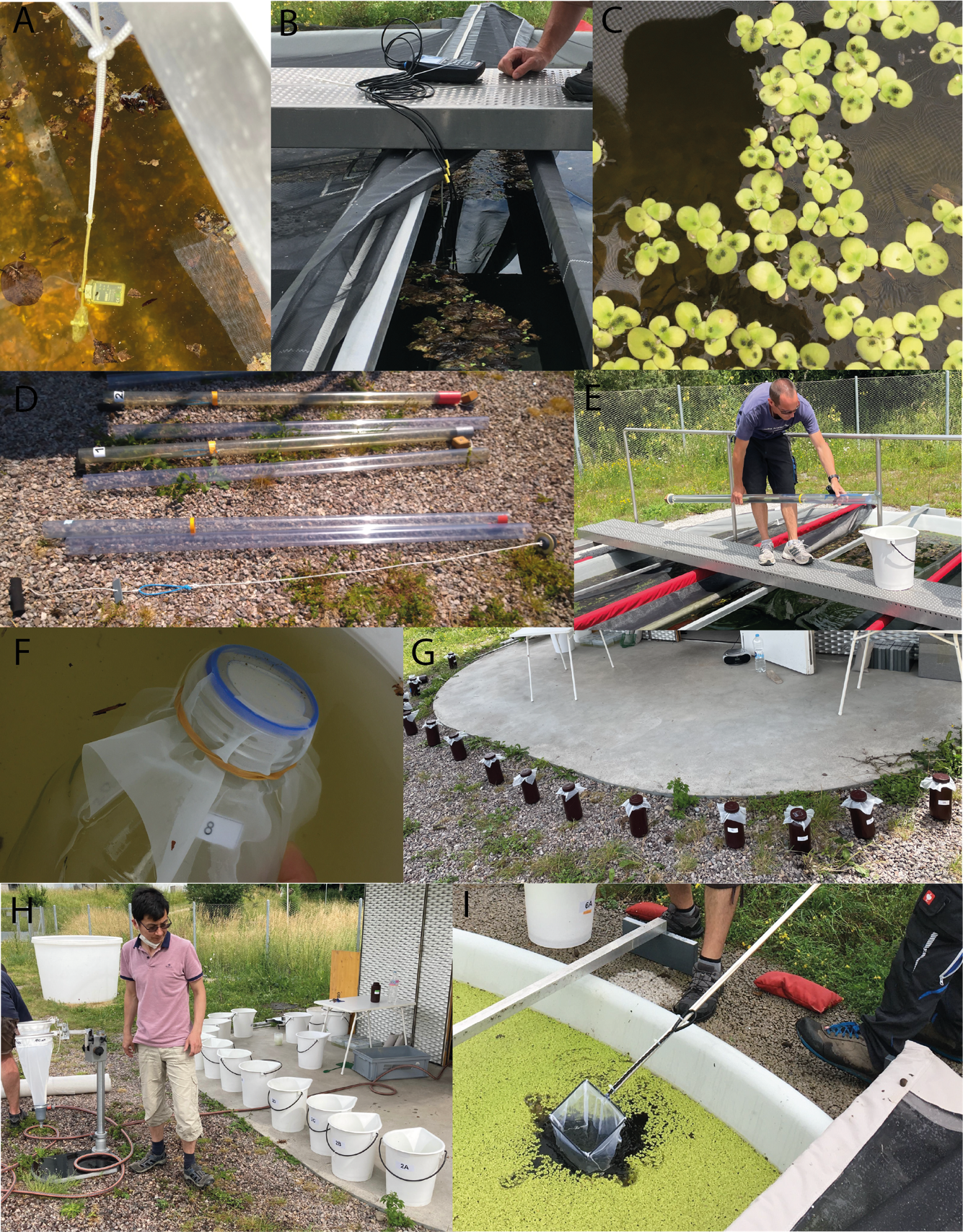

**Fig. S23 Data collection and sampling procedure** (A) Data loggers for the continuous recording of temperature and light data. (B) Dipping sonde for measuring the oxygen, pH and conductivity. (C) an exemplary picture that was used for estimating aphid densities. (D, E) Leibold-samplers that were used for the collection of water columns. Subsampling of the water used for (F) nutrient analysis, as well as (G) chlorophyll and phytoplankton analysis. (H) Buckets with the collected water and, on the left, the filtering equipment (mesh funnels) to collect the zooplankton. (I) Collection of *D. magna*.

**Table S1.** **Genes contain the SNPs that showed significant allele frequency differences between control and herbivory ponds.** *P*-values were determined using the beta-binomial mixed-effects model (from R-package glmmTMB v1.1.9 ^41^) with sampling time and pond block as the random factor. The three SNPs that are shown in Fig. 3 are highlighted in bold.

| **CHR** | **POS** | **P-value** | **GeneID** | **Putative function** |
| --- | --- | --- | --- | --- |
| CH1_R | 3296837 | 3e-09 | g3733 | - |
| CH1_R | 3297877 | 3.9e-08 | g3732 | - |
| CH1_R | 3318453 | 3.1e-09 | Daphnia_magna_D_magna_CH1_R_000761 | Cuticlin-1 |
| CH1_R | 3326249 | 4.8e-08 | FUN_002123 | - |
| CH1_R | 3328507 | 3.4e-08 | g3737 | Ribosome-binding protein 1 |
| CH1_R | 3331142 | 1.2e-08 | g3737 | Ribosome-binding protein 1 |
| CH1_R | 3331191 | 1.5e-08 | g3737 | Ribosome-binding protein 1 |
| CH1_R | 3345507 | 2.9e-08 | g3739 | - |
| CH1_R | 3345518 | 2.2e-08 | g3739 | - |
| CH1_R | 3349243 | 3.6e-08 | g3739 | - |
| CH2_L | 2021948 | 1.3e-08 | Daphnia_magna_D_magna_CH2_L_001270 | LIM/homeobox protein Lhx2 |
| CH2_L | 2233628 | 2.2e-09 | g4458 | Tachykinin-like peptides receptor 99D |
| CH2_L | 2233630 | 2.3e-09 | g4458 | Tachykinin-like peptides receptor 99D |
| CH2_L | 2233633 | 1.1e-09 | g4458 | Tachykinin-like peptides receptor 99D |
| CH2_L | 4270484 | 8.9e-10 | Daphnia_magna_D_magna_CH2_L_001057 | - |
| CH2_L | 4272659 | 2.4e-09 | Daphnia_magna_D_magna_CH2_L_001057 | Plasma kallikrein |
| CH2_L | 4281419 | 2e-09 | g4850 | Zwei Ig domain protein zig-8 |
| CH2_L | 4281563 | 2.8e-09 | g4850 | Zwei Ig domain protein zig-8 |
| CH2_L | 4281572 | 3.4e-09 | g4850 | Zwei Ig domain protein zig-8 |
| CH2_L | 4281868 | 1.4e-09 | g4850 | Zwei Ig domain protein zig-8 |
| CH2_L | 4282099 | 1.9e-09 | g4850 | Zwei Ig domain protein zig-8 |
| CH2_L | 4289074 | 4.9e-08 | g4850 | Zwei Ig domain protein zig-8 |
| CH2_L | 4289319 | 2e-08 | g4850 | Zwei Ig domain protein zig-8 |
| CH2_L | 4289322 | 6.7e-09 | g4850 | Zwei Ig domain protein zig-8 |
| CH2_L | 4291606 | 1.4e-08 | g4851 | - |
| CH2_L | 4315771 | 5.2e-08 | D_magna_XINB302771 | - |
| CH2_L | 4342533 | 1.3e-08 | g4857 | - |
| CH2_L | 4360738 | 3.9e-10 | D_magna_XINB302782 | Scavenger receptor class B member 1 |
| CH2_L | 4361537 | 2.1e-08 | D_magna_XINB302782 | Scavenger receptor class B member 1 |
| CH2_L | 4361829 | 2.9e-08 | D_magna_XINB302782 | Scavenger receptor class B member 1 |
| CH2_L | 4361831 | 4e-08 | D_magna_XINB302782 | Scavenger receptor class B member 1 |
| CH2_L | 4363525 | 2.7e-08 | g4861 | Scavenger receptor class B member 1 |
| CH2_L | 4371234 | 3.4e-09 | g4865 | Ribosome quality control complex subunit TCF25 |
| CH2_L | 4389279 | 1.1e-12 | Daphnia_magna_D_magna_CH2_L_000302 | - |
| CH2_L | 4401148 | 6.1e-09 | FUN_003244 | Luciferin sulfotransferase |
| CH2_L | 4411487 | 5.3e-08 | g4871 | Guanine nucleotide-binding protein G(o) subunit alpha |
| CH2_L | 4508578 | 3.5e-08 | D_magna_XINB302813 | Fructose-1,6-bisphosphatase 1 |
| CH2_L | 4508586 | 1.2e-08 | D_magna_XINB302813 | Fructose-1,6-bisphosphatase 1 |
| **CH2_L** | **4508591** | 1.5e-09 | **D_magna_XINB302813** | **Fructose-1,6-bisphosphatase 1** |
| CH2_L | 4508598 | 3.9e-09 | D_magna_XINB302813 | Fructose-1,6-bisphosphatase 1 |
| CH2_L | 4508660 | 2.2e-08 | D_magna_XINB302813 | Fructose-1,6-bisphosphatase 1 |
| CH2_L | 4508668 | 2.4e-08 | D_magna_XINB302813 | Fructose-1,6-bisphosphatase 1 |
| CH2_L | 4523214 | 3.6e-09 | g4890 | - |
| CH2_L | 4523278 | 1.8e-08 | g4890 | - |
| CH2_L | 4526022 | 3.7e-08 | D_magna_XINB302816 | - |
| CH2_L | 4527166 | 1.5e-09 | g4892 | - |
| CH2_L | 5844097 | 5.5e-08 | Daphnia_magna_D_magna_CH2_L_000939 | Neurofilament medium polypeptide |
| CH2_L | 5844814 | 2.9e-08 | Daphnia_magna_D_magna_CH2_L_000939 | Neurofilament medium polypeptide |
| CH2_L | 5844847 | 2.9e-08 | Daphnia_magna_D_magna_CH2_L_000939 | Neurofilament medium polypeptide |
| CH2_L | 5844848 | 5.6e-08 | Daphnia_magna_D_magna_CH2_L_000939 | Neurofilament medium polypeptide |
| CH2_L | 5844892 | 2.3e-09 | Daphnia_magna_D_magna_CH2_L_000939 | Neurofilament medium polypeptide |
| CH2_L | 5859492 | 8.7e-09 | Daphnia_magna_D_magna_CH2_L_000934 | - |
| CH2_L | 5860627 | 4.4e-08 | g5089 | Protein obstructor-E |
| CH2_L | 5954936 | 8.2e-09 | g5109 | Cyanophycinase |
| CH2_L | 5964503 | 1e-08 | Daphnia_magna_D_magna_CH2_L_000918 | Urea transporter 2 |
| CH2_L | 6043839 | 1.6e-09 | Daphnia_magna_D_magna_CH2_L_000906 | Homeobox protein prospero |
| CH2_L | 6043920 | 5.1e-09 | Daphnia_magna_D_magna_CH2_L_000906 | - |
| CH2_L | 6043969 | 3.1e-08 | Daphnia_magna_D_magna_CH2_L_000906 | - |
| CH2_L | 6044337 | 4.6e-10 | Daphnia_magna_D_magna_CH2_L_000906 | - |
| CH2_R | 2356529 | 5.9e-09 | g6043 | - |
| CH2_R | 2614664 | 1.2e-08 | g6071 | Transcription factor GATA-4 |
| CH3_L | 5712577 | 3.5e-08 | g7823 | - |
| CH4_L | 1865544 | 2.1e-15 | FUN_008680 | Histone H4 |
| CH5_L | 2622924 | 6.2e-12 | g12186 | - |
| CH5_L | 4457635 | 5e-08 | g12564 | Protein trachealess |
| **CH5_L** | **7535757** | 4.3e-09 | **Daphnia_magna_D_magna_CH5_L_000629** | **Ankyrin repeat domain-containing protein SOWAHC** |
| CH5_L | 7541932 | 1.8e-11 | g13107 | - |
| CH5_L | 7543997 | 2.7e-09 | D_magna_XINB310443 | - |
| CH5_L | 7544021 | 2.8e-08 | D_magna_XINB310443 | - |
| CH5_L | 7548886 | 4.7e-08 | FUN_012241 | - |
| CH5_L | 7548914 | 3.2e-09 | FUN_012241 | - |
| CH5_L | 7548927 | 2e-08 | FUN_012241 | - |
| CH5_L | 7549175 | 3.5e-08 | FUN_012241 | - |
| CH5_L | 7549238 | 5.4e-09 | FUN_012241 | - |
| CH5_L | 7553410 | 3.1e-08 | g13108 | N6-adenosine-methyltransferase TMT1A |
| CH5_L | 7649296 | 3.4e-08 | g13127 | - |
| CH5_L | 7649334 | 3e-09 | g13127 | - |
| CH5_L | 7649349 | 1.1e-08 | g13127 | - |
| CH6 | 8635704 | 3.4e-09 | D_magna_XINB301668 | Tetraspanin-3 |
| CH6 | 8650501 | 1.3e-08 | g15669 | Kinesin-like protein KIF3A |
| CH7_L | 1927186 | 3.1e-08 | g16459 | Serine/threonine-protein kinase N |
| CH7_R | 306771 | 2.9e-08 | g17612 | - |
| CH7_R | 312466 | 2.2e-08 | g17615 | - |
| CH7_R | 450106 | 3.4e-08 | Daphnia_magna_D_magna_CH7_R_000089 | - |
| CH7_R | 450870 | 2e-08 | g17671 | - |
| CH7_R | 450882 | 1e-09 | g17671 | - |
| CH7_R | 455399 | 6.9e-10 | g17673 | - |
| CH7_R | 455400 | 8e-10 | g17673 | - |
| CH7_R | 460581 | 3e-09 | Daphnia_magna_D_magna_CH7_R_000821 | - |
| CH7_R | 461672 | 4.6e-08 | g17675 | Tetraspanin-2A |
| **CH7_R** | **461798** | 2.7e-13 | **g17675** | **Tetraspanin-2A** |
| CH7_R | 461995 | 4.5e-09 | g17675 | Tetraspanin-2A |
| CH7_R | 462165 | 8.2e-12 | g17675 | Tetraspanin-2A |
| CH7_R | 462174 | 2.3e-12 | g17675 | Tetraspanin-2A |
| CH7_R | 462178 | 4.4e-13 | g17675 | Tetraspanin-2A |
| CH7_R | 462191 | 3.5e-12 | g17675 | Tetraspanin-2A |
| CH7_R | 462202 | 3.3e-10 | g17675 | Tetraspanin-2A |
| CH10_L | 146278 | 1.6e-09 | D_magna_XINB321143 | - |
| CH10_L | 147497 | 3.2e-08 | g43 | - |
| CH10_L | 147500 | 5.7e-08 | g43 | - |
| CH10_L | 147501 | 3.1e-08 | g43 | - |
| CH10_L | 151992 | 3.6e-08 | g44 | Histone deacetylase 6 |
| **CH10_L** | **152191** | **5.2e-08** | **g44** | **Histone deacetylase 6** |
| CH10_L | 2610115 | 4e-08 | g546 | Serine/threonine-protein kinase 26 |
| CH10_L | 974213 | 2.8e-10 | g263 | - |
